# GUIDE deconstructs genetic architectures using association studies

**DOI:** 10.1101/2024.05.03.592285

**Authors:** Daniel Lazarev, Grant Chau, Alex Bloemendal, Claire Churchhouse, Benjamin M Neale

**Affiliations:** Department of Mathematics, Massachusetts Institute of Technology, Cambridge, Massachusetts, USA; Renaissance School of Medicine at Stony Brook University, Stony Brook, New York, USA; Stanley Center for Psychiatric Research, Broad Institute of MIT and Harvard, Cambridge, Massachusetts, USA; Analytic and Translational Genetics Unit, Massachusetts General Hospital, Boston, Massachusetts, USA; Program in Medical and Population Genetics, Broad Institute of MIT and Harvard, Cambridge, Massachusetts, USA; Novo Nordisk Foundation Center for Genomic Mechanisms of Disease, Broad Institute of MIT and Harvard, Cambridge, Massachusetts, USA

## Abstract

Genome-wide association studies have revealed that the genetic architecture of most complex traits is characterized by a large number of distinct effects scattered across the genome. Functional enrichment analyses help provide some biological interpretation of associated variants but more work is needed to further translate GWAS hits into meaningful biological insights. Thus, we set out to leverage the genetic association results from many traits with a view to identifying the set of modules, or latent factors, that mediate these associations. The identification of such modules may aid in disease classification as well as the elucidation of complex disease mechanisms. We propose a method, Genetic Unmixing by Independent Decomposition (GUIDE), to estimate a set of statistically independent latent factors that best express the patterns of association across many traits. The resulting latent factors not only have desirable mathematical properties, such as sparsity and a higher variance explained for the latent factors that are significantly associated with a given trait, but are also able to single out and prioritize key biological features or pathophysiological mechanisms underlying a given trait or disease. Moreover, we show that these latent factors can isolate biological pathways as well as epidemiological and environmental influences that compose the genetic architecture of complex traits.

---

Mappings between the human genome and phenome—already complicated by the high dimensional structures involved—are further convoluted by polygenic and pleiotropic relationships. Deconvolving the genetic architecture underlying a given set of phenotypes may be achieved by learning a set of modules, or latent factors, which mediate between a set of genetic variants and a set of traits. These latent factors can be viewed as the components of the genetic architecture of the outcome traits and may relate to biochemical mechanisms (*e*.*g*., LDL cholesterol and its impact on coronary artery disease) or genetic influences on exposures (*e*.*g*., smoking as a risk factor for lung cancer). By using genetic association results across a wide range of traits it may be possible to identify different subsets of genetic variants that point to such biochemical or exposure related processes and in turn facilitate the interpretation of results from genome-wide association studies (GWAS). This is especially crucial for complex traits—those emerging from the interplay of several intertwined molecular and environmental mechanisms represented by distinct latent factors—for which the number of causal variants can be in the tens of thousands [1].

Existing methods that group subsets of genetic variants, such as genomic structural equation modeling (genomic SEM) [2], mash [3], and pleiotropic decomposition regression (PDR) [4], are not necessarily informed by the traits they affect, cannot be applied to a large number of traits, or require user-specified model parameters, architecture or latent factors.

One approach that addresses some of these concerns, FactorVAE, uses a variational autoencoder (VAE) to learn a latent representation of the data that seeks to ‘disentangle’ it by finding independent factors [5]. In a similar vein, a recently developed method, Decomposition of Genetic Associations (DeGAs) [6, 7], applies truncated singular value decomposition (TSVD) to GWAS summary statistics to derive a set of latent factors for this purpose. Applying singular value decomposition (SVD) to a matrix of summary statistics produces a diagonal matrix containing the singular values that is left- and right-multiplied by orthogonal matrices. Truncating these two orthogonal matrices based on a user-specified number of latent factors gives a set of orthonormal vectors that—viewing this arrangement as a three-layer network (Fig. 1)—may be considered as the normalized weights from a layer of single nucleotide polymorphisms (SNPs) to a new layer of latent factors and from the latter to a layer of traits. Applications of DeGAs have already included, for example, newer measures of polygenic risk scores that are informed by these factors [8].

**Figure 1:**
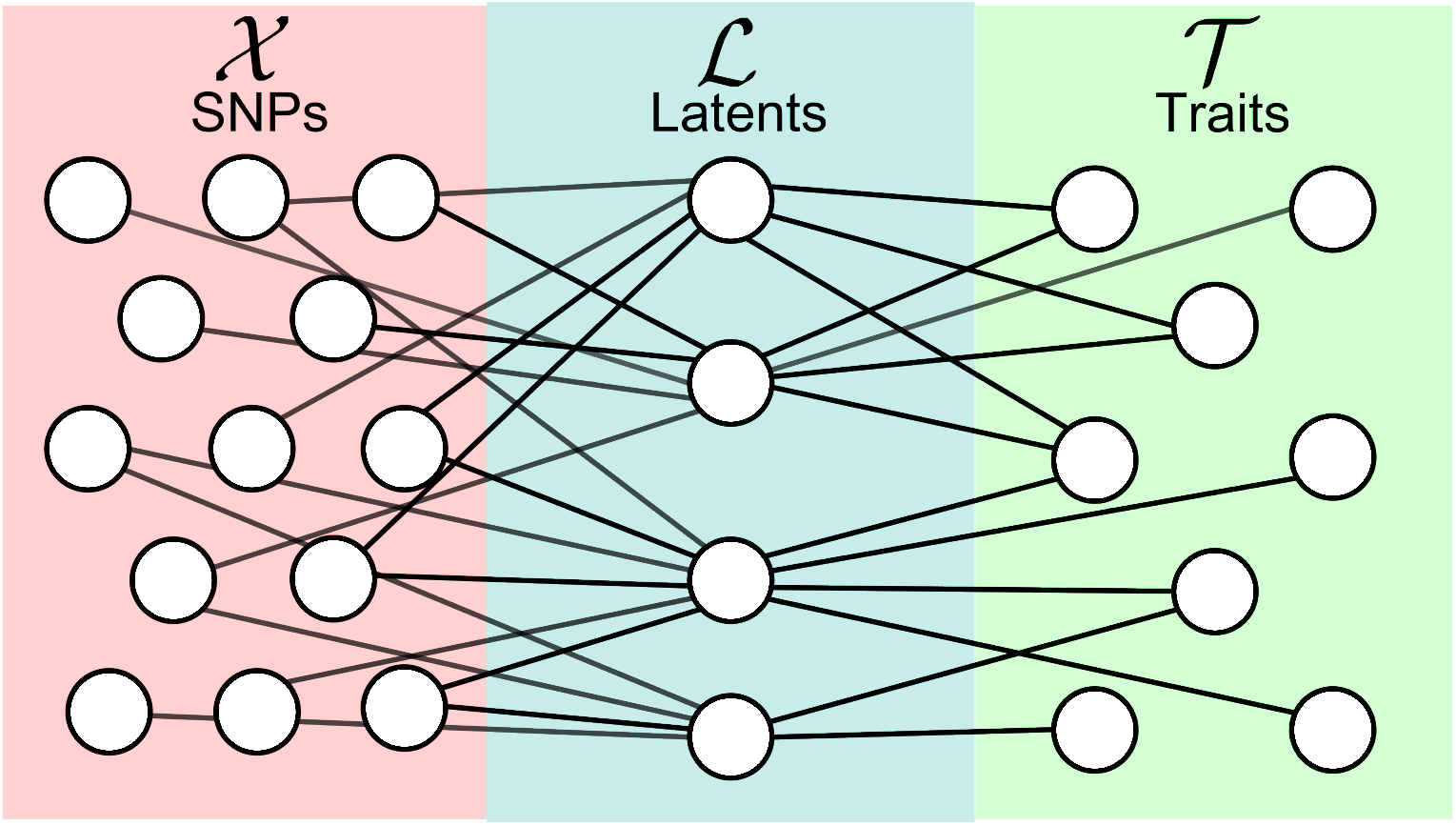
An example of a three-layer SNP-phenotype network. Here, 𝒳 is the layer of SNPs, ℒ the layer of latent factors, and 𝒯 the layer of traits. We write, *M* := |𝒳 |, *L* := |ℒ|, *T* := |𝒯 |.

A shortcoming of DeGAs, however, stems from the fact that the SVD solution is one of an infinite number of possible solutions to this decomposition problem: one can multiply by any orthogonal matrix and its inverse (the transpose in this case) to get another possible decomposition. In other words, starting from any possible decomposition of the summary statistics, this is essentially the problem of choosing a basis for the latent factors that gives a set of maximally sparse, independent, and interpretable factors.

Independent Component Analysis (ICA) is a statistical method for decomposing a mixed signal into a set of statistically independent ones (*e*.*g*., separating individual conversations from a recording of multiple conversations in the famous cocktail party problem) [9, 10, 11, 12]. It has been successfully used in many applications in genetics and genomics, for example, as a denoising or dimensionality reduction alternative to principle components analysis (PCA) [13]; to decompose *Escherichia coli* transcriptome data into regulatory or mechanistic modules governing key aspects of the bacterium’s biological activity, such as its virulence or response to oxidative stress or antibiotics [14, 15]; and combined with other methods such as canonical correlation analysis [16] or more recently, PCA [17, 18], to similarly decompose neuroimaging genetics data consisting of separate SNP and functional magnetic resonance imaging (fMRI) data, or GWAS summary statistics of brain image-derived phenotypes, respectively.

Our method, Genetic Unmixing by Independent Decomposition (GUIDE), uses TSVD and a two-sided application of ICA—simultaneously for both the SNPs-to-latents and latents-to-traits weights—to determine a set of sparse and independent latent factors given GWAS summary statistics. Unlike previous approaches, GUIDE also includes an additional method, also based on ICA, for estimating the number of latent factors and a method for estimating the significance of the learned weights by comparing them with other possible models of the same data (Methods).

The latent factors determined by GUIDE can illuminate relations between different diseases, help interpret results from the GWAS of any trait, and perhaps motivate a reclassification of diseases that more accurately reflects their interrelations via, for example, shared symptoms or common disease mechanisms. We apply GUIDE to several curated subsets of summary statistics from the UK Biobank, including those used in [6], allowing for a direct comparison with DeGAs. An interactive browser presenting the GUIDE model for this dataset can be found at guide-analysis.hail.is, allowing for the analysis of traits of choice from the UK Biobank using GUIDE latent factors.

Using GUIDE to analyze Alzheimer’s disease, for example, we show how GUIDE, driven solely by data without any user-specified modeling choices or parameters, prioritizes *APOE* and other nearby genes as contributing to a distinct mechanism leading to Alzheimer’s disease, distinguishing it from a general high cholesterol pathway that has long been associated with Alzheimer’s disease [19, 20]. Applying GUIDE to another complex trait, forced expiratory volume in one second (FEV1), showcases its ability to disentangle molecular from psychosocial or epidemiological effects on a given trait, identifying latent factors that influence FEV1 by affecting smoking behavior, cardiovascular health, immune or developmental processes, standing height, and thoracic cavity volume.

We provide a brief overview of GUIDE in Section 1.1 of Results (with full details in Methods), followed by a review of the salient properties of GUIDE latent factors and other features of our method in Section 1.2. Section 1.3 focuses on the GUIDE analysis of Alzheimer’s disease, while Section 1.4 is a deep dive into FEV1 and related traits.

## 1 Results

### 1.1 Overview and motivation for the GUIDE method

As described in full detail in Methods, GUIDE simultaneously applies Independent Component Analysis (ICA) to both the SNP and trait weights matrices corresponding to a given set of summary statistics after obtaining an initial decomposition of the summary statistics matrix using singular value decomposition (SVD). This is motivated by the following insights.

Entropy is a measure of randomness or uniformity of a probability distribution, and its value is lowered by the presence of information or constraints, which make some events more probable than others. Therefore, on a finite support (*e*.*g*., [0, 10]) the distribution with the highest entropy is the uniform distribution, while on an infinite support (*e*.*g*., the real number line, R), among distributions with a finite variance, the Gaussian distribution maximizes entropy [21]. Solving a sixty-year-old problem posed by Claude Shannon, it was shown [22] that normalized sums of independent random variables have a higher entropy with every random variable added, eventually converging to the maximum entropy value as the sum becomes normally distributed in the limit. In other words, these sums are not only Gaussian in the limit, per the central limit theorem, but are also ‘more Gaussian’, in the sense of having a higher entropy, at every step.

The GUIDE model assumes the existence of independent latent modules mediating the effects of genetic variants on traits (see Fig. 1). Since the effects of these latent factors are summed over, the factors themselves are hidden in the data, and may seem normally distributed due to the above considerations. The solution, therefore, would be not only to decompose the summary statistics using SVD, but also to find a basis for the network weights that *minimizes* the entropy—one that is maximally non-Gaussian—in order to recover the original latent factors. Luckily, ICA does exactly that. ICA is built on the idea of maximizing non-Gaussianity via the minimization of entropy (or, equally, maximization of negentropy). Although entropic measures are the optimal cost functions from a mathematical perspective [22, 9, 10], in practice, ICA implementations use computationally favorable proxies of entropy, such as kurtosis or some notions of sparsity, which do not require the estimation of a probability distribution function at every step.

The implementation of ICA we use, FastICA, is based on a particular set of optimal cost functions derived using the maximum entropy principle that were found to outperform previous proxies based on sparsity or kurtosis [11, 12].

Nevertheless, these older proxies are still useful in assessing the performance of our solutions (SI, Sec. 1.2) and in gaining intuition for the kinds of solutions that GUIDE finds. More specifically, as we will see in the next section, since we apply ICA to both the 𝒳 → ℒ and 𝒯 → ℒ weights, the weight matrices to and from the GUIDE latent factors are maximally sparse and kurtotic, that is, with few, sharp peaks of signal that capture most of the interlayer relationships of a given latent factor with the relevant SNPs and traits.

An outline of the GUIDE method can be found in Algorithm 1 in Methods.

### 1.2 Properties of GUIDE and a Novel Measure of Weight Significance

Compared with other methods that have used ICA, *e*.*g*., [17, 18, 16, 14, 15], there are several important differences that define and distinguish GUIDE. These include: (a) an underlying model architecture in which ICA is used to find a basis for both 𝒳 → ℒ and ℒ → 𝒯 weights simultaneously, thus having the latent factors informed by both genetic variants and traits; (b) a novel method for estimating the number of latent factors required for a given dataset as well as a new heuristic based on the entropies of different models; and (c) a novel method for estimating a “weight significance value”, or *w*-value, for every 𝒳 → ℒ or ℒ → 𝒯 weight, allowing for quantitative measures of significance of the GUIDE model weights and automated exploratory analyses using user-specified significance thresholds. As detailed in Methods (Sec. 3.5), *w*-values are computed by comparing a given weight against the distribution of the corresponding weights from all other possible models decomposing the given summary statistics matrix. In particular, the *w*-value is the probability of randomly drawing the given weight value or a more extreme one given the distribution of corresponding weights from all possible models. Notably, though similar in spirit, *w*-values are not *p*-values because there is no specific null hypothesis weight against which a given model’s weight is compared, but rather the infinite set of other possible latent models of dimension *L*.

In summary, GUIDE facilitates the deconvolution of biological effects and molecular mechanisms, enabled by the model’s architecture and the particular application of ICA, allowing it to learn statistically independent intermediate modules mediating the effects of variants on traits and grouping traits based on shared mechanisms or effects.

For direct comparison with DeGAs, we used the ‘all’ dataset consisting of 235,907 variants and 2,138 phenotypes from [6], which originated from the UK Biobank (UKB). We will only refer to DeGAs as such when applying it to the ‘all’ dataset on which it was tested by the authors in [6] and for the particular analyses they had used it for; otherwise, especially in subsequent sections using new curated subsets of the UK Biobank or novel analyses, we will write ‘TSVD’.

As the analyses in the next sections will detail, GUIDE not only nominates groupings of candidate SNPs for further experimental validation, but (perhaps more importantly) disentangles the pathways or mechanisms mediating the effects of genetic variants on traits of interest, even for related traits. For example, a quick review of a GUIDE model of the UKB ‘all’ dataset revealed two latent factors, one affecting adipose tissue traits and fat mass (*e*.*g*., ‘Trunk total mass’, ‘Arm fat mass (right)’, and ‘VAT (visceral adipose tissue) volume’) and the other affecting lean mass (*e*.*g*., ‘Arm lean mass (left)’,’Total fat-free mass’, ‘Trunk lean mass’). These latent factors implicated not only SNPs near genes involved in the physiology common to both sets of traits, such as *FTO*, but also others involved in pathways specific to each one, for example, *NPY4R, RIN3*, and *GCKR* for the lean mass latent factor, and *PPAR-γ, ADRB3*, and 16p12.3 (upstream of *GPRC5B70*) for the fat mass latent factor [23, 24].

As shown in Figs. 2(a) and 3, GUIDE latent factors explain the variance in the traits using far fewer factors than those of TSVD and FactorVAE, confirming GUIDE’s ability to pick a basis for the latent factors that sparsifies the weights from the latent factors to both the SNPs and the traits (see also Figs. S4 and S5). Put differently, the top GUIDE latent factors always explain more of the variance than their TSVD counterparts. This effect is also qualitatively seen in Figs. S8 and S9, which explicitly compare the SNPs-to-latents and the latents-to-traits weight matrices for GUIDE and DeGAs, showing that, in both cases, the GUIDE matrices are sparser with much more concentrated signal as compared with the DeGAs matrices, for which the weight is more uniformly spread across the latent factors.

**Figure 2:**
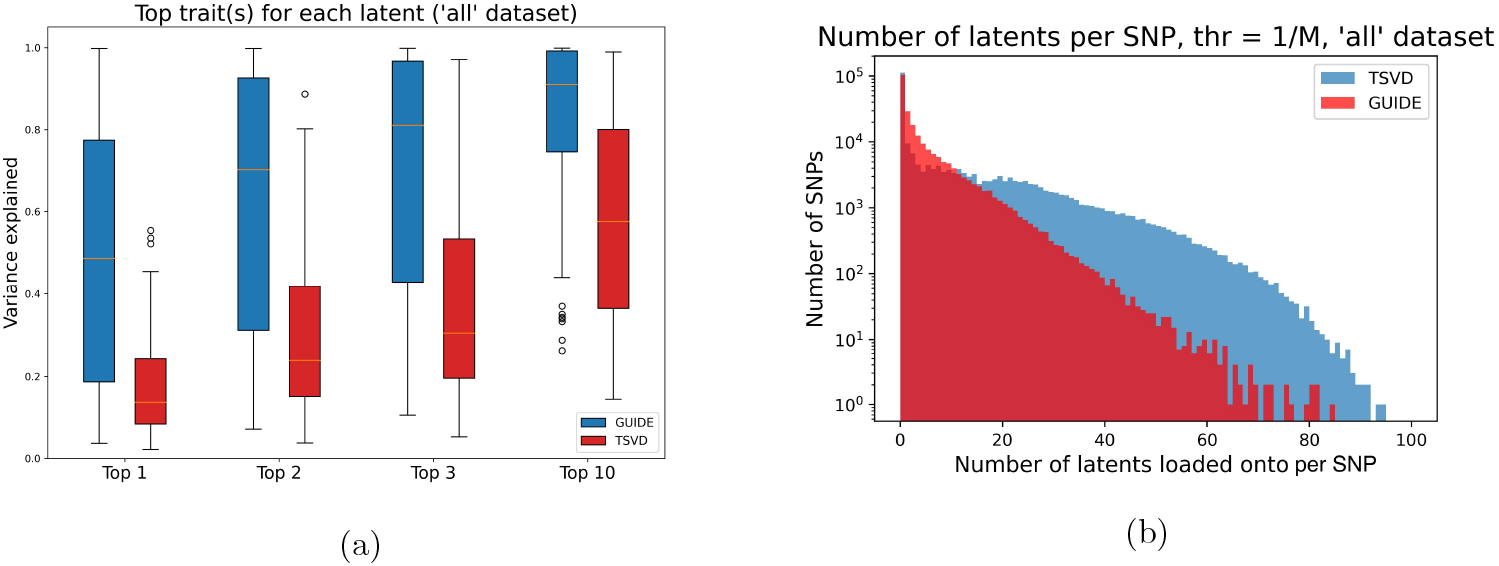
(a) Contributions of the top 1, 2, 3, and 10 traits to the normalized variance explained by each latent factor for both GUIDE and TSVD model weights. See Fig. S8 for the variance explained curves for all 100 latent factors for GUIDE and for TSVD. Center line, median; box limits, upper and lower quartiles; whiskers, 1.5x interquartile range; points, outliers. (b) Number of latent factors each SNP loads on to with a squared weight greater than 1*/M*. SNPs load onto fewer GUIDE factors compared to TSVD factors.

Fig. 3 compares the fraction of the genetic variance components (see Methods for the definition) by the top three latent factors of GUIDE, DeGAs, and FactorVAE, showing that for the top factors of each model, GUIDE latent factors explain a larger fraction of the genetic contribution to the given traits using the same underlying set of SNPs.

**Figure 3:**
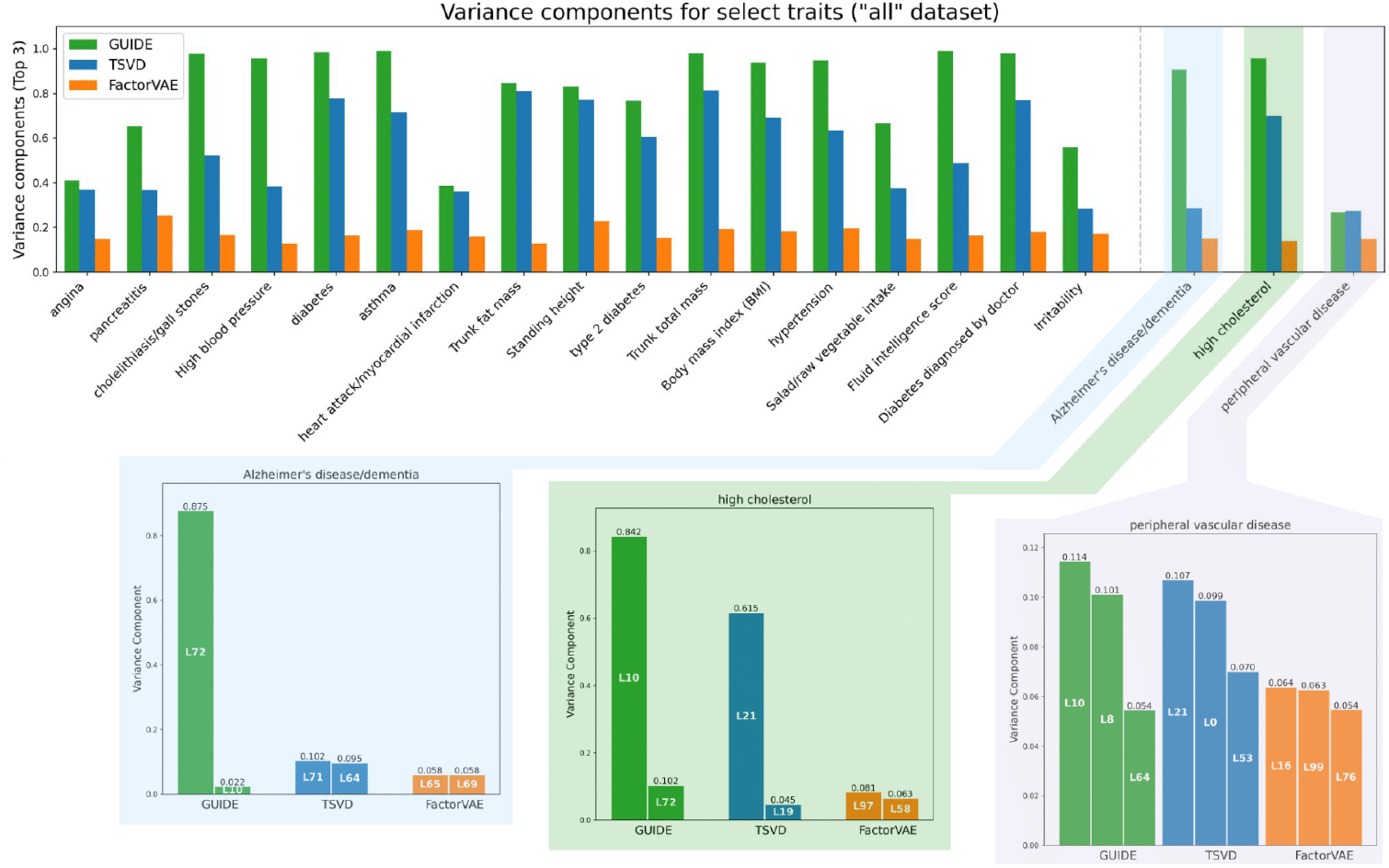
Variance components for select trait using the ‘all’ dataset from [6]. We report the sum of the scores for the top three latent factors for each trait.

To more specifically characterize the biology of certain traits, we applied GUIDE to eight subsets of the UK Biobank consisting of SNPs significant for a given trait (*e*.*g*., ‘High Blood Pressure’, as in Fig. 4) with *p* < 5 ×10^−8^ and with all other traits correlated to the given trait with *r*^2^ > 0.5 (see Table S3 in SI). Figs. S6 and S7 show the GUIDE and TSVD ℒ → 𝒯 heat maps for models with *L* = 100 and *L* = 10, respectively, for the high blood pressure subset in Table S3. While the *L* = 100 GUIDE model significantly out-performed the corresponding TSVD model in terms of both sparsity and concentration of signal, the difference was not significant for the *L* = 10 models. Motivated by these observations, and by the fact that the qualitative discrimination of models afforded by these types of heat maps is not as precise and will not be feasible for vastly larger values of *M* or *T*, we further examined more quantitative measures of GUIDE’s performance as a function of the number of latent factors, *L*.

**Figure 4:**
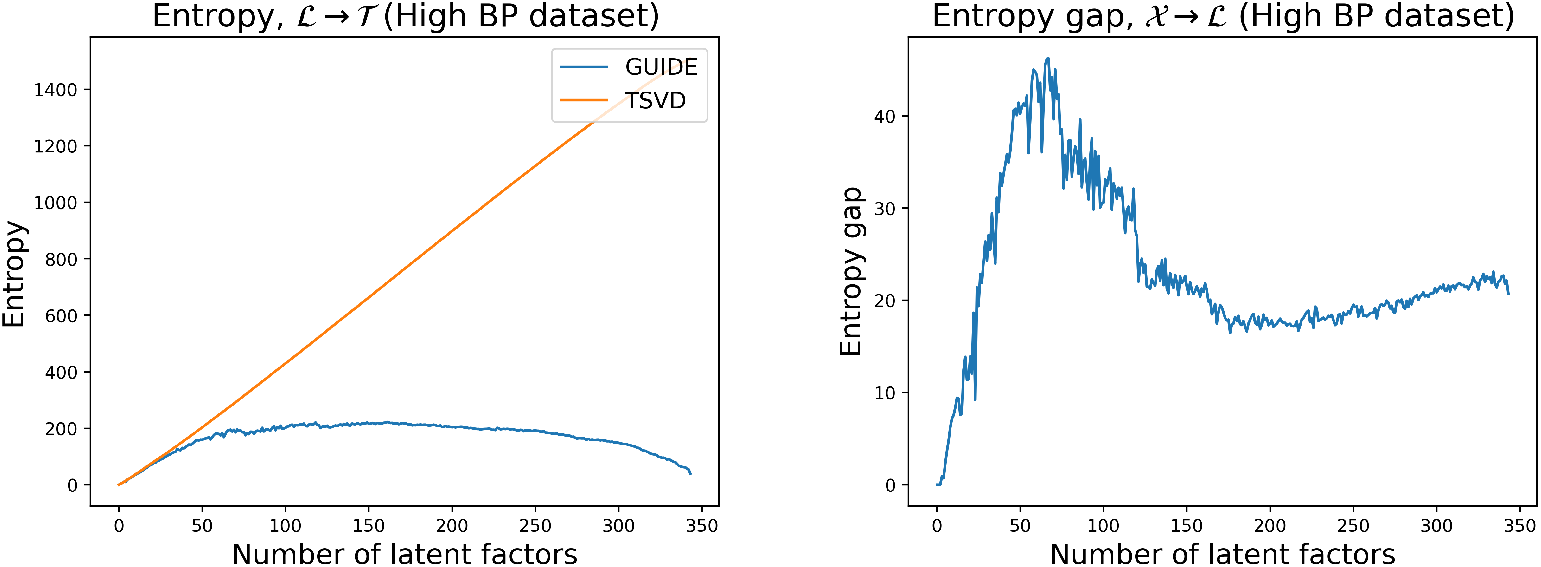
Plots of the entropy for the ℒ → 𝒯 weights and entropy gap for the 𝒳 → ℒ weights (defined as the difference between the TSVD model entropy and the GUIDE model entropy) as a function of the number of latent factors in the each model for the High Blood Pressure dataset, defined in Table S3, consisting of 1183 SNPs significant for high blood pressure (*p* < 5 × 10^−8^) and 344 traits that are correlated to it with *r*^2^ > 0.5. The entropy plot for the 𝒳 → ℒ weights and the entropy gap plot for the ℒ → 𝒯 can be found in the Supplementary Information.

While, as mentioned above, using entropy is computationally expensive in the execution of the ICA portion of our method, it can still be used ex post facto to compare the performance of GUIDE relative to TSVD. We note that a lower entropy corresponds to a more informative signal. We also look at the entropy gap, defined as the difference between the TSVD model’s entropy and the GUIDE model’s entropy, with the understanding that the higher the entropy gap, the more information is gained by GUIDE relative to the TSVD model.

The first plot in Fig. 4 compares the entropies for the TSVD and GUIDE models as a function of the number of latent factors in each model, while the second shows the entropy gap, defined as the difference between the TSVD model’s entropy and the GUIDE model’s entropy, also as a function of the number of latent factors. Fig. 4 makes it clear that the difference in the entropies is negligible for *L* = 10 and is quite significant for *L* = 100, which explains our previous qualitative observation comparing GUIDE’s performance between Fig. S6 and Fig. S7.

Moreover, referring to the ℒ → 𝒯 entropy plot for the ‘High BP’ dataset (see Table S3 in the SI) in Fig. 4, we see that the entropy plateaus at around *L ≈* 50, suggesting that GUIDE models with at least that number of latent traits should capture most of the information contained in the traits of that dataset given the same dataset’s SNPs. The 𝒳 → ℒ entropy gap in Fig. 4 shows that the largest entropy gap is for *L* ∈ [40, 70], further suggesting that a choice of *L* in this range would allow for more information gain (for the 𝒳 → ℒ weights) and a more discernible advantage for GUIDE. (Referring again to the entropy plot in Fig. 4, it is clear that the corresponding entropy gap would be monotone increasing, but that this would be driven purely by the increasing entropy of the TSVD model, which would nonetheless mean that for larger value of *L*, the GUIDE weights would be consistently sparse and concentrated.) Indeed, using our model selection algorithm to estimate the optimal number of latent factors (Methods), we obtained a value of 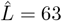 for the ‘High BP’ dataset. Figure S13 shows the ℒ → 𝒯 entropy plot for the UKB dataset.

We also prepared a subset of the UK Biobank multi-ancestry PanUKB dataset (pan.ukbb.broadinstitute.org) by filtering to only include genotyped SNPs, traits with a *z*-score greater than 4, and excluding missing regression coefficients, consisting of 635,964 SNPs and 677 traits, which we will refer to as the ‘UKB *Z* > 4 dataset’. We built an *L* = 100 GUIDE model for this dataset.

Using the contribution scores between the ℒ and 𝒯 layers, and a threshold 1*/T*, which is the value corresponding to a completely uniform distribution of the weights (Methods), we defined the sets of traits for each latent factor with squared weights greater than 1*/T* as an indication of which traits are significantly represented by each latent factor.

For the UKB *Z* > 4 dataset, we found that there are 13.94 traits per GUIDE latent on average, compared to 67.38 traits per TSVD latent, even though GUIDE latent factors load onto 68% of all traits with greater than uniform weight, compared to 54% for the TSVD latent factors. Thus, not only are the GUIDE latent factors more sparse (with almost 5 times less traits per latent), but they nevertheless account for more of the traits than TSVD.

For the 𝒳 and ℒ layers, we used a 1*/M* threshold for the same reasons. Fig. 2(b) shows the number of latent factors each SNP loads onto with squared *W*_*XL*_ weights greater than 1*/M*, confirming GUIDE’s ability to deconvolve shared SNP effects by sparsifying interlayer weights, as will be discussed in more detail in the next section.

Reproducibility and generalizability of latent factors represent a particularly difficult problem and an active area of research in the representation learning community [25, 26, 27]. To be clear, these are two distinct but related properties: (1) generalizability: a model’s ability to consistently perform well across different datasets, not only the dataset it was initially trained or tested on; and (2) reproducibility: finding latent factors that are consistently reproduced across different datasets.

To assess generalizability, we constructed a GUIDE model on an LD-pruned set of FinnGen data [28] (*r*^2^ < 0.2, with 188,140 SNPs and 2408 traits). We computed the *w*-values for the weights for the FinnGen and UKB models to quantify the performance by measuring, for example, the percentage of traits with maximum weights that pass a significance threshold of *p* < 10^−8^. These and other statistics are summarized in Table 1.

**Table 1:** Assessing performance of GUIDE models for UKB and FinnGen using *w*-values for ℒ → 𝒯 weights.

| Dataset | % traits with max weight $-\log(w) > 8$ | mean of max $-\log(w)$ | std. dev. of max $-\log(w)$ | max $-\log(w)$ over all traits |
| --- | --- | --- | --- | --- |
| UKB | 29.3 | 30.7 | 25.2 | 125 |
| FinnGen | 47.0 | 14.5 | 7.65 | 67.5 |

In the next section, we confirm these results qualitatively in the case of Alzheimer’s disease, showing GUIDE’s ability to yield biologically meaningful insights across two different datasets, despite various significant confounding effects, such as LD, genetic correlation, and stratification, which GUIDE does not account for.

Nevertheless, to determine potential reproducibility of GUIDE latent factors, we isolated the 121 traits that are shared between the UKB and FinnGen datasets. As there were so few SNPs shared between the UKB and FinnGen datasets, we retained all variants in each set for this comparison (namely, 235,907 for UKB and 188,140 for FinnGen). Building GUIDE models for both datasets we found several latent factors that matched based on the overlap in the traits that significantly load onto them. These include a factor, found in both UKB and FinnGen GUIDE models, which significantly loads onto ulcerative colitis and aortic dissection [29, 30]; another that significantly loads onto Alzheimer’s disease, anxiety disorder, constipation [31], and pneumonia [32]; and lastly, a factor that significantly loads onto prostate cancer, inguinal hernia, polycystic ovarian syndrome, and multiple myeloma [33, 34, 35, 36].

As we stress in the Discussion below, while this comparison provides a somewhat limited and qualitative assessment of reproducibility, the observed recurrence of these latent factors serves to demonstrate that reproducibility is feasible, even across very different datasets and in the presence of confounding factors. We hope that future methodological advances that take into account these confounding properties would show improved reproducibility.

### 1.3 Isolating molecular pathophysiological pathways of Alzheimer’s disease

The analyses in this section and the next use the GUIDE model built using the ‘all’ dataset described in Sec. 1.2.

In order to characterize and contextualize each of the top latent factors for Alzheimer’s disease (AD), we also considered the traits, “high cholesterol” and “peripheral vascular disease,” for which GUIDE latent factors revealed common pathophysiological mechanisms.

For each of these traits we identify the top latent factors based on their *genetic variance components*, defined as a normalized sum of 𝒳 → ℒ or ℒ → 𝒯 weights (Methods). We use Genomic Regions Enrichment of Annotations Tool (GREAT) [37] to qualitatively assess the interpretability of these latent factors and explore the top genes that are enriched for them. GREAT estimates the functional significance of *cis* regulatory regions, and is also able to account for distal binding sites. Though GREAT does not seem to directly account for LD, this is not an issue in our case given that we already LD pruned all the datasets we used. GREAT does, however, account for the length of gene regulatory domains.

Fig. 3 highlights the top two or three latent factors for Alzheimer’s disease, high cholesterol, and peripheral vascular disease for GUIDE, TSVD, and FactorVAE together with their genetic variance components. Fig. 5 presents the top traits and loci contribution scores for the GUIDE and TSVD latent factors for AD and high cholesterol. To improve interpretability, we used ANNOVAR [38] to assign the top 1-3 genes for each locus implicated by our method in Figure 5 and in Figure 6 in the next section, though there is some uncertainty in the assignment of some of the genes (*e*.*g*., for loci with several nearby genes). Fig. S15 reports the *p*-values and binomial fold enrichments given by GREAT for the shared GO processes of GUIDE latent 72 and TSVD latent 71.

**Figure 5:**
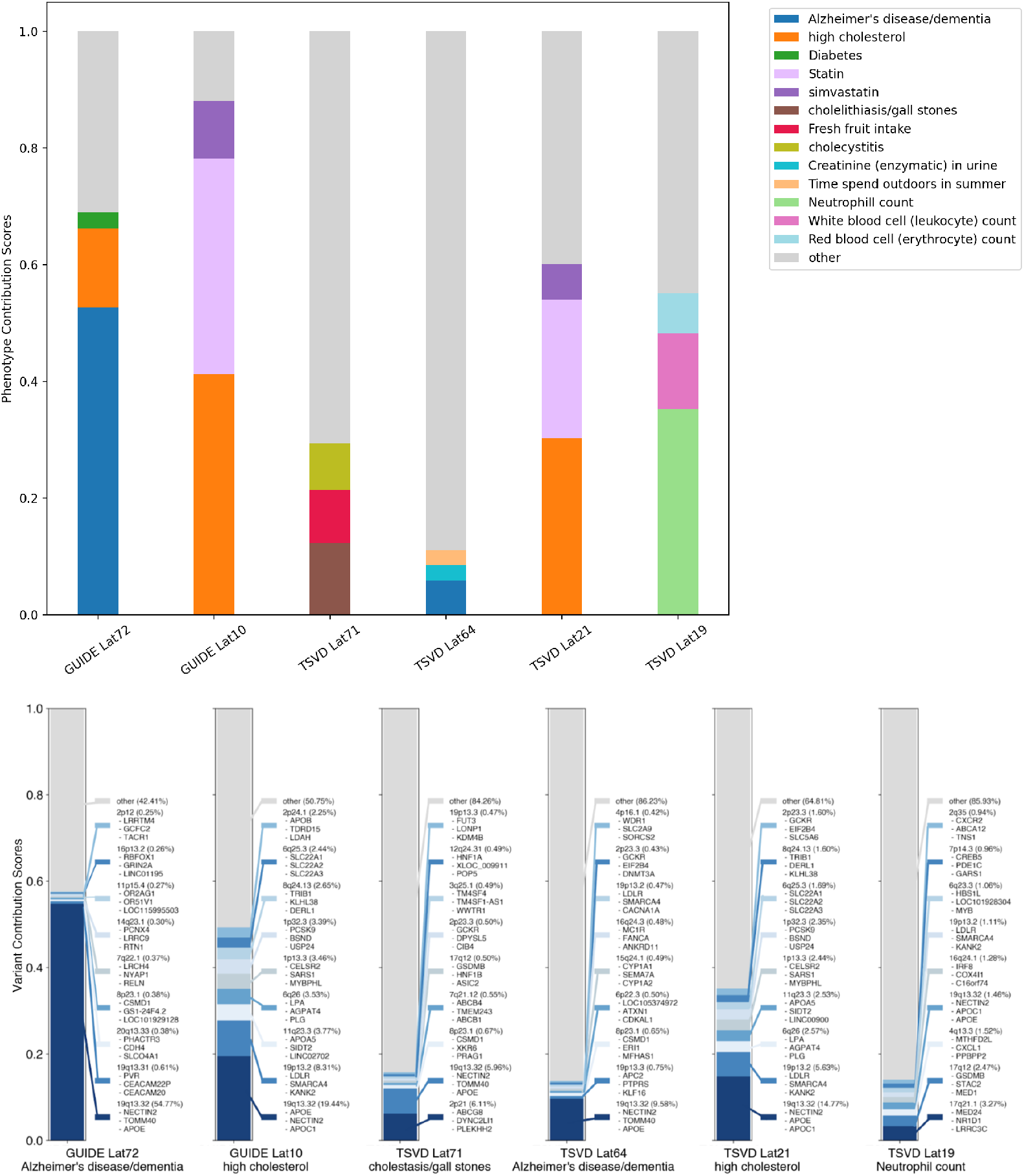
Top three contributions from traits and top nine contributions from loci onto the relevant GUIDE and TSVD latent factors as defined in Fig. 3. We determined the genes for the top SNPs using ANNOVAR [38]. The corresponding contribution scores for each locus are in parentheses. The contribution score due to each gene within a given locus is based on the ANNOVAR annotations.

**Figure 6:**
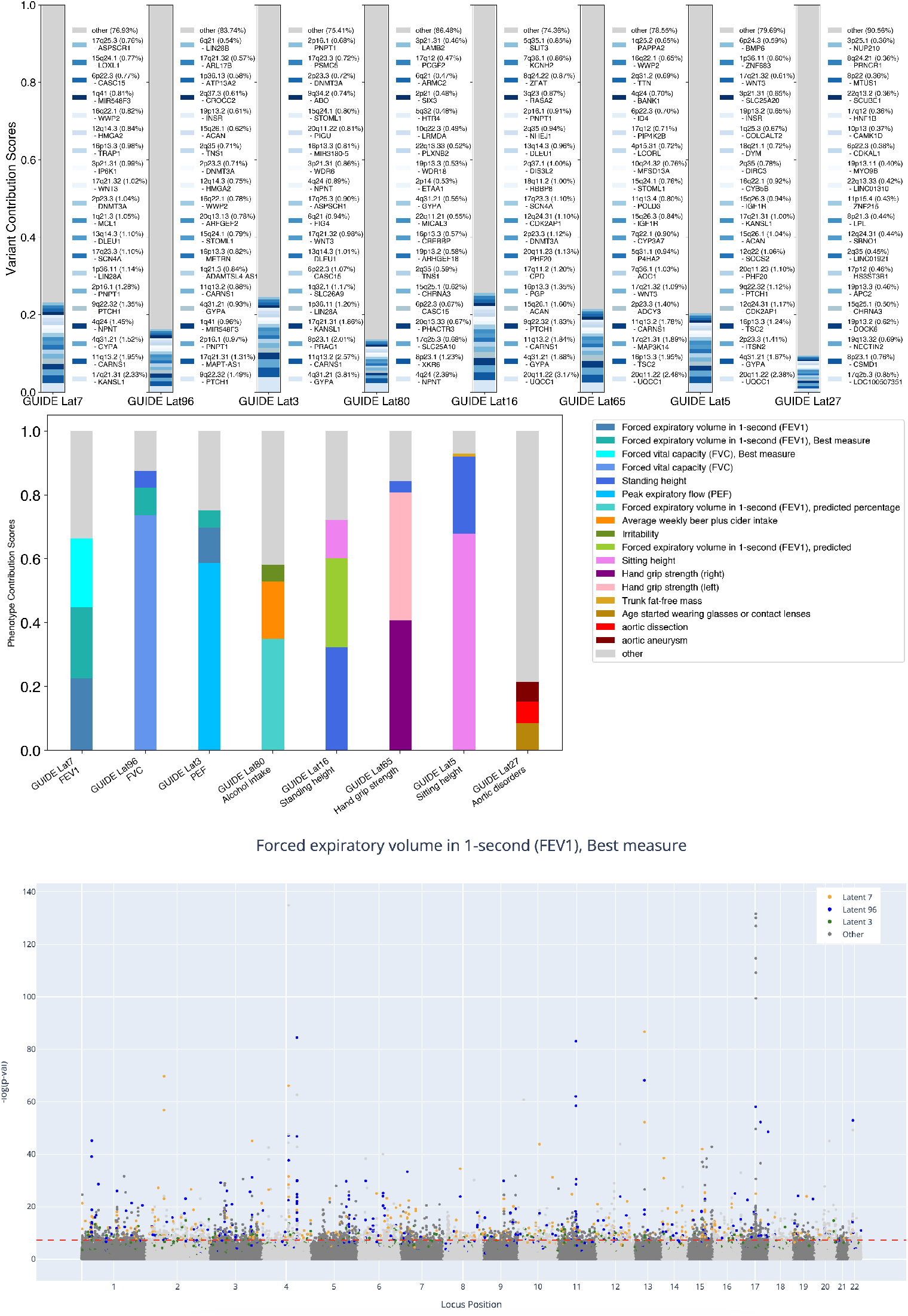
Top latent factors for FEV1 traits, showing the top variants’ (top panel) and phenotypes’ (middle panel) contributions to each latent factor. The latent factors are ordered in descending order by their contribution to the trait, “FEV1, Best measure”. Bottom panel: Manhattan plot highlighting the variants tagged by the top three latent factors contributing to FEV1 traits. The significance threshold, given by the dotted red line, has been set at *p* = 5 × 10^−8^.

For Alzheimer’s disease, GUIDE latent 72, which has a variance contribution score of 87.5% (see subsection 3.3 in Methods), is highly enriched for the *APOE* gene, with 49 of the top 54 SNPs in the 19:45020859-45728059 region near that gene having a contribution score of 50.9% [39]. Moreover, GUIDE prioritizes loci on chromosome 16 near *RBFOX1* and on chromosome 8 near *CSMD1*, which have only recently been implicated in AD [40, 41], as well as loci such as 20q13.33, which have previously been found to be associated with AD [42].

On the other hand, GUIDE latent 10 loads on other cholesterol genes, such as *APOA, APOB, APOC, LDLR, LPA, LPL, LPLR*, and *PCSK9* [43], and can thus be interpreted as mediating the effects of genes involved in cholesterol metabolism more generally (even though it also loads onto *APOE*, but less significantly than latent 72). GUIDE latent 72 can then be understood as mediating disease processes involving *APOE*, but not as part of the cholesterol metabolic pathway per se. Some of the genes implicated by latent 10, such as *SORT1* (chr. 1) [44], have only recently been found to be significant for AD.

In contrast, TSVD latent 71, the top TSVD factor for Alzheimer’s disease with a variance contribution score of 10.2%, mostly picks up a host of related genes such as *APOC, APOB, LDLR*, among others, without significant enrichment for *APOE*, and with the top 50 loci—nonspecifically spanning chromosomes 1-4, 7-15, 17, 19, and 20—having a contribution score of only 12.7%. Furthermore, TSVD latent 71 significantly implicates GO processes that are not exclusively related to Alzheimer’s pathophysiology, such as “positive regulation by host of viral process” (*p* = 4.8 × 10^−8^), “drug transport” (*p* = 1.16 × 10^−7^), and “spleen development” (*p* = 2.8 × 10^−6^).

For high cholesterol, GUIDE latent 10 contributes a genetic variance component of 84.2%, and its high contribution is consistent with its earlier interpretation as a factor significant for a larger group of cholesterol metabolism genes. GUIDE latent 72, highly enriched for the *APOE* gene is the second most contributing factor, with a genetic variance component of 10.2%.

Unlike GUIDE, TSVD does not pick up on the connection with the cholesterol latent factors first identified in connection with Alzheimer’s disease, instead prioritizing two other, unrelated factors.

The case of peripheral vascular disease shows GUIDE’s usefulness even when its latent factors have low genetic variance component scores that are comparable with TSVD’s factors. The genetic variance components for the top three latent factors for GUIDE and TSVD are both around 27%, with each top factor (latent 10 for GUIDE and latent 21 for TSVD, Fig. 3) matching the top latent factor for high cholesterol. Nevertheless, while the second GUIDE latent (*L*8) prioritizes vascular and cardiac muscular processes, such as “myofibril assembly” (*p* = 3.97 × 10^−16^) and “sarcomere organization” (*p* = 4.13 ×10^−19^), and with a significant number of SNPs implicating genes such as *TTN, GJA1, SCN5A, WDR1, MYH7*, and *CSK*, among others [45, 46, 47, 48, 49, 50], the second TSVD latent (latent 0) does not seem to identify processes directly related to vascular disease pathophysiology, prioritizing GO biological processes such as “regulation of glucose metabolic process” (*p* = 1.03 × 10^−23^) and “neural plate pattern specification” (*p* = 2.87 × 10^−18^). And while TSVD latent 53, the third contributing factor to peripheral vascular disease, does include GO processes related to cardiovascular morphology and function, it only predominantly prioritizes *PITX2*, and none of the other important genes implicated by GUIDE latent 8. For GUIDE, the third highest factor, GUIDE latent 64, significantly prioritizes genes such as *AHR, PML, GDNF*, and *IRX1-3*, which were not tagged by TSVD latent factors, but were recently implicated in important mechanisms contributing to vascular disease, such as vascular inflammatory pathways, atherosclerosis, endothelial dysfunction, and oxidative stress [51, 52, 53, 54].

Running a GUIDE model on an LD-pruned set of FinnGen data [28] (*r*^2^ < 0.2, with 188,140 SNPs and 2408 traits) revealed additional insights not found in the UKB models. For example, the FinnGen model had one top latent for AD that predominantly implicated 14q24.2, the locus where *PSEN1* is located, another implicated the 9p21 locus associated with vascular dementia and AD [55].

Both *APOE* and *PSEN1* are known to be implicated in different clinical manifestations of Alzheimer’s disease, attesting to GUIDE’s ability to yield relevant and biologically insightful latent factors across two different datasets. Though not finding reproducible latent factors in this case, the above results do show GUIDE’s ability to generalize, that is, to produce significant and insightful results across different datasets (see previous section for further discussion).

Fig. S14 shows the percent shared SNPs for the top 500 SNPs associated with each latent factor in Fig. 3, measured using the Jaccard index (Methods). Among the GUIDE latent factors, there is only an overlap of 8.0% between latents 72 and 10, due to the *APOE* -associated SNPs in latent 10, as mentioned earlier. Among the TSVD latents, there is more overlap between factors 71, 64, 21, and 19, suggesting that a more significant part of the genetic variance component—as compared with the sparser GUIDE factors—is contributed by TSVD factors due to double counting of certain contributing SNPs, which inflates the score.

GUIDE latent 10 and TSVD latent 21, both of which correspond to nonspecific genes involved in cholesterol metabolism, share 73.4% of their top 500 SNPs. This kind of overlap is to be expected, especially for highly effective TSVD factors, since TSVD can, for some factors, pick out a basis that happens to closely align with that derived by GUIDE. TSVD latent 21 is quite effective given that, unlike the factor comparison with Alzheimer’s disease where none of the top TSVD latents were shared, we do see that latent 21 is again implicated in peripheral vascular disease, like its GUIDE counterpart, latent 10. Figure 5 summarizes some of the above discussion, showing the top contributing genes and traits for each of the relevant latent factors identified in Fig. 3.

All in all, Fig. S14 summarizes some of the trends we have seen in this section: GUIDE latent factors are more independent (in the sense of lower mutual information; Methods) and sparse (except where certain processes naturally overlap), allowing them to home in on a wider range of important genetic variants while differentiating between distinct biological processes and disease mechanisms.

### 1.4 Deconvolution of FEV1 reveals environmental and molecular factors

Whereas the last section showed GUIDE’s ability to deconvolve the molecular mechanisms contributing to a given trait, in this section we show, using the example of forced expiratory volume in one second (FEV1), GUIDE’s ability to disentangle and stratify the behavioral or environmental mechanisms from the molecular ones contributing to a particular end trait.

Fig. 6 shows the stacked plots for the top latent factors for FEV1 in descending order by the magnitude of their genetic variance components for FEV1. The top two panels show the breakdown of the contributions to each latent factor by loci and traits, respectively, while the bottom panel shows the Manhattan plot for ‘FEV1, best measure’, where the loci tagged by the top three latent factors are colored in.

Latent factor 7 contributes to FEV1 through immunological and epigenetic regulatory mechanisms, tagging genes such as *KANSL1, WNT3* and *GYPA* [56, 57]. *Latent factors 80 and 27 in Figure 6 (see also Fig. S12, Supplementary Information) are the top contributing factors for smoking behavior, implicating loci with genes such as CHRNA3*, which encodes the alpha 3 subunit of the nicotinic acetylcholine receptor, which is known to associate with smoking behaviors [58]. Interestingly, after building a GUIDE model using the same traits but removing traits related to smoking behavior, the corresponding latent factors—determined by matching contributions other traits and variants—in the new model do not implicate those same genes, as seen in Figure S15.

The phenotypic contributions to latent 65 are dominated by the left and right hand grip strength traits, while the variants tagged by latent 65 are associated with genes such as *TTN, UQCC1, TSC2, CARNS1*, and *IGF1R*, suggesting that this factor represents cardiovascular health and fitness as mechanisms affecting FEV1. Latent 5 contributes to FEV1 by way of developmental processes that lead to increased lung volume by virtue of having a larger thoracic cavity, implicating genes such as *UQCC1, PTCH1*, and *CDK2AP1* [59]. Unsurprisingly, latent 5 is the top contributing latent factor for sitting height and the third for standing height.

As in the last section, we also compared these results with the same GUIDE model on the LD-pruned set of FinnGen data [28] (*r*^2^ < 0.2, with 188,140 SNPs and 2408 traits). Given that the FinnGen trait set is completely different from that of the UKB—and, in particular, FinnGen does not have any FEV1 traits—we instead looked at all the traits related to respiratory disorder, such as asthma, chronic obstructive pulmonary disease (COPD), interstitial lung disease (ILD), or respiratory distress of the newborn. The top latent factors implicated for these traits loci such as 1p31.1 associated with lung cancer in never-smokers [60], 2q11.2 (which seems to be implicated in immune processes/cancer in the lungs) [61], and 2p24, which seems to be associated with FEV1, asthma, and COPD [62, 63, 64].

The lower panel in Fig. 6 shows a Manhattan plot for the trait “FEV1, best measure”, where we highlighted the variants tagged by the top three GUIDE latent factors contributing to this trait.

Let *g*_sig_ be the number of variants implicated by GUIDE that are above the significance threshold, *g*_total_ be the total number of variants implicated by GUIDE, *n*_sig_ be all variants above the significance threshold and *n*_total_ be the total number of variants. We can then define the enrichment of the top three GUIDE latent factors by,

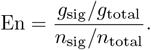

For the Manhattan plot in Fig. 6, En = 24.5, indicating, by a naive interpretation of this metric, that the top three GUIDE latent factors tag 24.5 times more significant variants as would be expected were variants picked at random. However, given that LD is nonuniformly spread across variants, with a preference for significant variants, this metric cannot be taken at face value. Conversely, stringent LD pruning—which would presumably fix this problem—may make it more difficult for GUIDE to find significant variants given that most of them would be removed. The most sensible approach thus seems to appreciate—qualitatively—that En > 1 indicates that the latent factors have picked up more significant variants than would be expected were they to be chosen randomly, but not to overinterpret the actual value of En (though perhaps allowing for comparisons).

## 2 Discussion

We developed GUIDE as a method for disentangling polygenicity and pleiotropy into more coherent subsets of genetic effects that might relate to specific pathways or exposures relevant to outcome traits under study. GUIDE uses summary statistics data from GWAS and PheWAS to build a sparse and interpretable three-layer network consisting of SNPs, traits, and an intermediate layer of latent factors. GUIDE not only uses a simultaneous two-sided application of ICA to find the most optimal basis (informed by both the genetic and phenotypic data) for these latent factors, but also estimates the number of required latent factors using an approach that, unlike current model selection methods, works better when applied to larger, noisier data sets. Moreover, we developed a method to estimate the *w*-values of all GUIDE model weights (compared to a null distribution of all possible alternative model weights; Methods), allowing for a principled quantitative assessment of the model’s performance and reliability.

The independence of the latent factors and the sparsity of the weights connecting them with the SNP layer 𝒳, on one side, and the trait layer 𝒯, on the other, are not only mathematically and computationally convenient properties. More importantly, as we have shown, these properties make the GUIDE latent factors more biologically interpretable and allow them to extract more information from the underlying genetic architecture. For example, as seen in the application to Alzheimer’s disease in Section 1.3, GUIDE latent factors were able to separate out some of the main genetic contributions to the disease due to the *APOE* genes, thus discriminating between the pathophysiological pathways due to these genes versus other ones in the cholesterol metabolic pathway, which were captured by another latent factor.

It is important to note, however, that the success of the ICA portion of the GUIDE algorithm rests on the assumption that there exist discrete modules mediating between the SNPs and traits (see Fig. 1). This assumption fails, for example, for genetic variants with infinitesimal effects on given traits, which induce a continuous variance-covariance structure that cannot be written as a sum of discrete non-Gaussian modules. Here, TSVD would be the better approach. In fact, using the approach outlined in Methods (3.9) GUIDE can be used to build models that capitalize on this phenomenon by extracting the discrete modules (the latent factors) as before, but using the remaining Gaussian components (which ICA cannot distinguish) to analyze the continuous contributions to a trait’s variance.

Moreover, the results of GUIDE are influenced by the traits included in the data, as we showed in Sec.1.4 in the analysis of FEV1 traits using GUIDE models with and without smoking traits. Naturally, since GUIDE latent factors are informed by the given genetic variants and traits, larger datasets produce better results, suggesting that broader and better powered systematic GWAS would help build better informed GUIDE models. Though we focused on diagnoses rather than course, outcome or treatment response traits in this paper, we hope that using larger sets of traits that include the latter will also allow GUIDE to glean new insights that will improve our understanding of the dynamics of human illnesses.

Of course, the expectation that using larger datasets would better inform GUIDE latent factors is mitigated by the presence of linkage disequilibrium (LD) and genetic covariance.

We have shown how the use of model weight *w*-values could be used to quantify the performance of, and thus assess, GUIDE’s ability to generalize across datasets in the sense of producing reliable and biologically meaningful results. Nevertheless, the problem of finding latent factors that are reproduced in different datasets—thus representing biological pathways or processes that transcend the particular populations from which the data were derived—remains a largely open problem. We report reassuring results where we found, using subsets of UKB and FinnGen summary statistics with matching traits, particular latent factors that loaded onto the same traits with significant weight strengths.

Given that GUIDE does not explicitly account for LD or genetic covariance, the latent factors it derives are biased by the presence of both, although to a somewhat lesser degree than TSVD. For example, in Fig. S10, we plotted the genetic variance components for the 2,692,064 million SNPs by 4,357 traits UKB dataset GUIDE model with *L* = 200 (as estimated by our model selection algorithm, Methods). We noticed that for this set, the genetic variance components for the top three TSVD latent factors had a score close to 1 for nearly all traits, while the top three GUIDE latent factors had much lower scores (even lower than for the curated subsets of the UK Biobank used in Results). This is likely due to large LD and genetic covariance inflating the weights of the TSVD latent factors (which, as mentioned above in Sec. 1.3, usually have a much larger number of above-average-loading traits per latent factor) to a larger extent than the sparser GUIDE weights, an effect we have observed in Fig. S14. We confirmed this effect in simulations, where we observed that LD inflated the estimated model *w*-values, with significantly greater inflation for TSVD models than for GUIDE models (see Methods and Fig. S18). LD pruning in these simulations did minimize these effects, especially for GUIDE.

Though these problems can be partially avoided by LD and genetic covariance pruning, and running a GWAS of many traits from the same cohorts, as opposed to using GWAS from different studies, as mentioned in Sec. 1.4, overly aggressive LD pruning may hinder GUIDE’s ability to find significant variants since most of them would be removed. The definitive solution, which we hope to pursue in future work, therefore seems to require explicitly incorporating the effects of both LD and genetic covariance, allowing for models that accurately account for both intra- and inter-layer effects. This would allow using the information contained in these intra-layer relationships to inform the model instead of biasing its results. Such models also hold promise to implicate causal variants, serving as a new avenue for finemapping. By directly accounting for LD and genetic covariance, while also perhaps incorporating newer information-maximizing algorithms [65], we hope that future iterations of our method will be even better equipped to learn biologically-relevant latent modules, while also accounting for population differences and ancestry.

In summary, GUIDE does provide a principled way to decompose genetic associations for complex traits into more coherent subunits. The interpretation of the factors that emerge from a GUIDE analysis is facilitated by the strength of the relationship to the outcome traits as well as the biological annotation of the genetic variants driving the signal. These resulting factors may facilitate the interpretation of genetic analyses and suggest groups of SNP effects to pursue together for functional follow-up studies.

## 3 Methods

### 3.1 Preliminaries

In our network model, we start with a set of SNPs, 𝒳, and a set of traits, 𝒯. Then the matrix of estimated regression coefficients (“betas”), 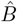, gives the weights of the edges connecting every SNP-trait pair. Our goal is to determine the weights to and from a layer of intermediate latent factors, ℒ, that best represent and summarize the relationships between 𝒳 and 𝒯. In other words, our basic problem is to find a decomposition,

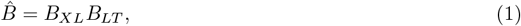

where *B*_*XL*_ is the *M* × *L* matrix of 𝒳 to ℒ weights and *B*_*LT*_ is the *L* × *T* matrix of ℒ to 𝒯 weights, with *L* < *T* < *M*. Intuitively, the ‘true’ latent layer ℒ is an intermediate, lower dimensional space that maximally extracts the genetic information encoded in 𝒳 and is thus a better, more biologically informed representation of the resulting traits in 𝒯.

Let *u*_*𝓁*_ be the *𝓁*th column vector of *B*_*XL*_, and 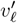 be the *𝓁*th row vector of *B*_*LT*_ :

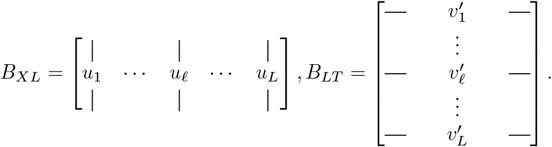

Then we can write (1) as the sum

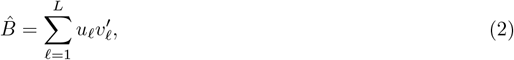

where, in the context of our network model, each product of vectors gives the matrix of weights to and from a single latent factor.

Therefore, the number of latents traits *L* is, in theory, equal to the rank of 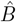:

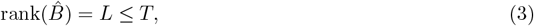

where the inequality here is due to the property that the rank of a matrix is at most equal to the least of its dimensions (in our case 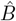 is a *M T* matrix, with *T* < *M*). In reality, however, most datasets are sufficiently noisy so that rank 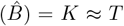. Therefore, more sophisticated methods are needed to estimate *L* (see Sec. 3.9).

Our task is essentially to decompose 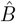 as in Eq. (1). Of course, if *B*_*XL*_, *B*_*LT*_ satisfy (1) then so would

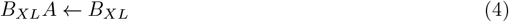

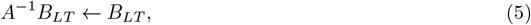

since

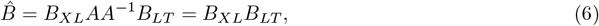

where *A* ∈ GL(*L*, ℝ) is a transformation matrix—a member of the family of all *L* × *L* invertible matrices with entries in ℝ—that chooses the proper scaling and orientation of the weights. (Of course, for *L* = 1, *A* and *A*^−1^ are simply multiplicative constants.) By whitening 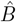 we instead look for *A* ∈ O(*L*, ℝ), that is for an *L* × *L* orthogonal matrix.

Our solution to this decomposition problem, GUIDE, will be outlined in the next section. We note here that we can easily extend GUIDE to allow one to include predefined latent factors in the latent layers by defining a *discrepancy matrix*, 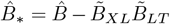, where the second term includes the weights to and from the predefined set of latent factors. Starting from 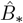 we then derive the rest of the latent factors to fill out the network using our method as before.

### 3.2 An optimal basis for the latent factors

We start by using truncated singular value decomposition (TSVD) to obtain an initial decomposition of 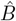:

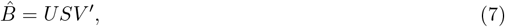

where *S* is an *L* × *L* diagonal matrix consisting of the nonzero eigenvalues of 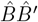, *U* is an *M* × *L* orthogonal matrix consisting of the eigenvectors of 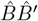 corresponding to the (squared) eigenvalues in *S*, and *V* is an *T* × *L* orthogonal matrix consisting of the same for 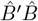. In the next section we address how to estimate the appropriate value for *L* using our method.

Let *C*_*n*_ be the *n* × *n* centering matrix, an orthogonal projection given by

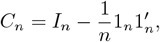

where *I*_*n*_ is the *n* × *n* identity matrix and 1_*n*_ is the vector of all ones. Even though the nonzero entries in 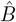 may be drawn from a distribution with a mean of zero, the sampled values will not be exactly mean-centered. Thus, we first mean center both the rows and columns of 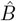 by applying left- and right-multiplying by the appropriate centering matrix:

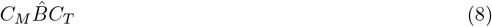

Define the *L* × (*M* + *T*) matrix by concatenating *U* ^′^ and *V* ^′^:

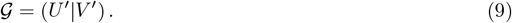

As a consequence of the centering, *U* and *V* are mean-centered as well. Then, using the fact that *A*^−1^ = *A*^′^, we may write (4) and (5) as

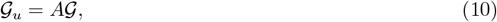

where 𝒢_*u*_ is the *L* × (*M* + *T*) unmixed matrix each of whose rows consists of the *M* + *T* weights to and from a latent trait. Written this way, it is clear that we are looking for the unmixing matrix *A* that would pick a basis for the latent factors that is simultaneously informed by both the SNPs in 𝒳 and the traits in 𝒯. The weight matrices *W*_*XL*_, *W*_*LT*_ to and from the latent factors in ℒ are then given by (modulo scaling of the weights),

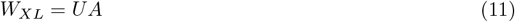

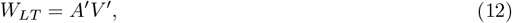

where we solve for *A* by applying Independent Component Analysis (ICA)to (10) using the FastICA algorithm [12, 9]. Note that 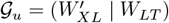.

With this in mind, we rewrite (6), now in the new basis, as

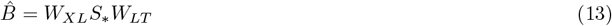

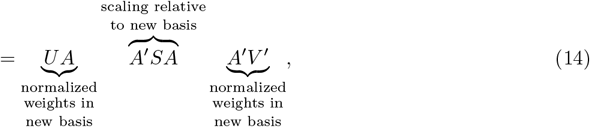

where

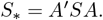

Of course, *S*_*_ is not diagonal, which reflects the fact that the components of the new basis are generally not orthogonal in the original (prewhitened) feature space (if they were, then *A* would be a permutation matrix, and *S*_*_ would be diagonal with elements equal *S* but reordered). For example, let 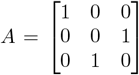, the permutation matrix that swaps the second and third components, and write 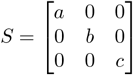. Then 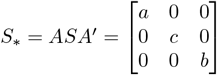. As this simple example illustrates, *S*_*_ is nothing more than a scaling relative to the new basis. While a permutation keeps the mutual orthogonality of the components intact, a rotation generally does not, which may provide some intuition for the existence of nonzero off-diagonal entries in *S*_*_ when *A* is a rotation matrix.

Also note that 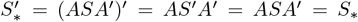. In the whitened feature space, the ICA components are orthogonal, and in that space *A* is a permutation matrix.

Alternatively, we can view the decomposition as

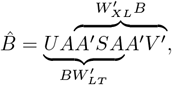

which is further addressed in Section 3.4

The method detailed in this section is summarized below in Algorithm 1.

To check GUIDE’s ability to recover discrete additive components, we conducted a simulation with 10^6^ variants and 1000 traits, with 100 independent latent factors and added noise. Fig. 7 shows how weights estimated using GUIDE can recover the real (simulated) weights modulo scaling for both 𝒳 → ℒ and ℒ → 𝒯 weights, while using TSVD alone fails to do so.

**Figure 7:**
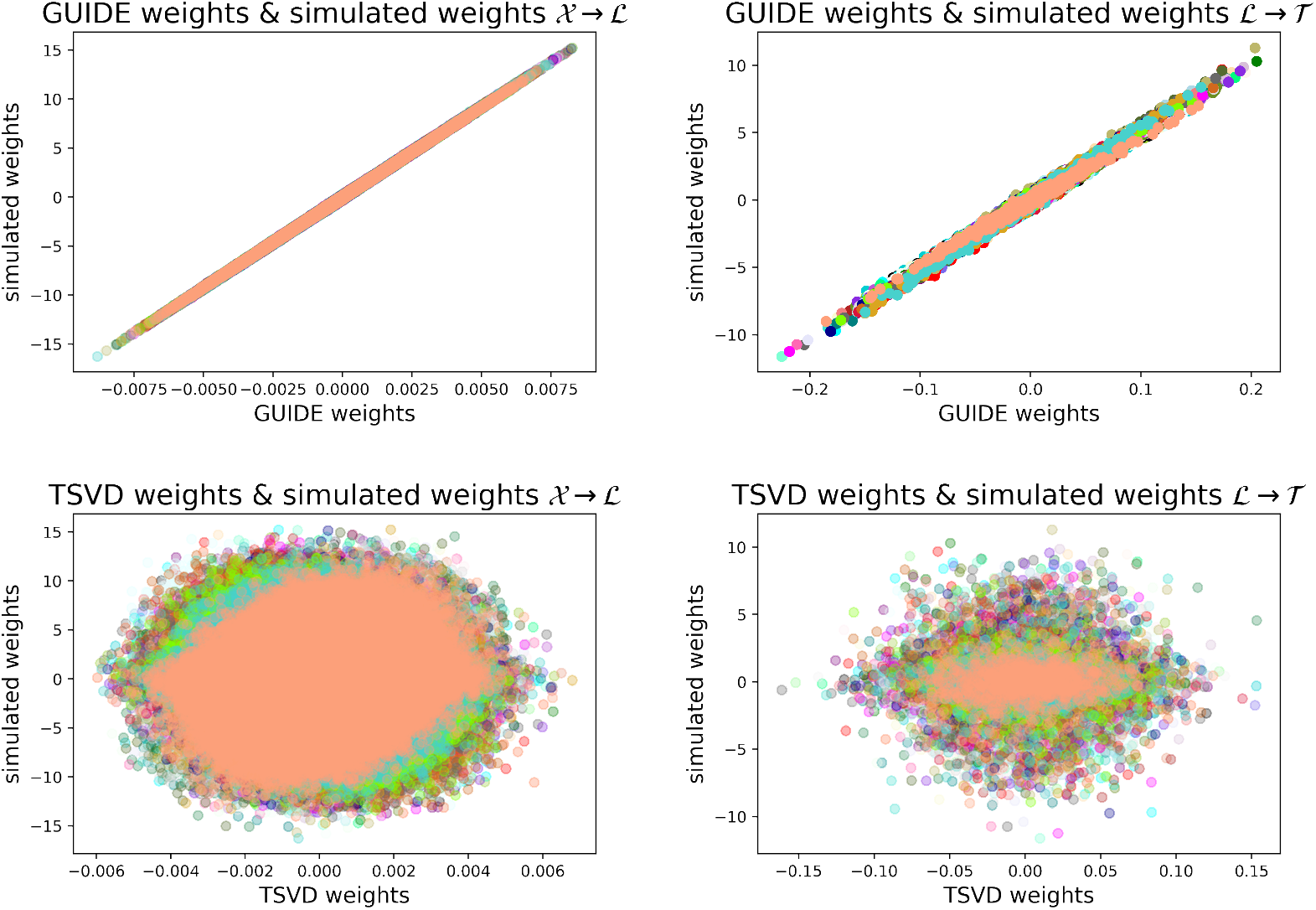
Scatter plot comparing the real (simulated) weights with the estimated ones using GUIDE versus TSVD. In this simulation, we used *M* = 10^6^, *T* = 1000 and *L* = 50, and corrected for the permutation and sign differences between the real and estimated latent factors. Each color corresponds to a different latent factor.

#### Algorithm 1

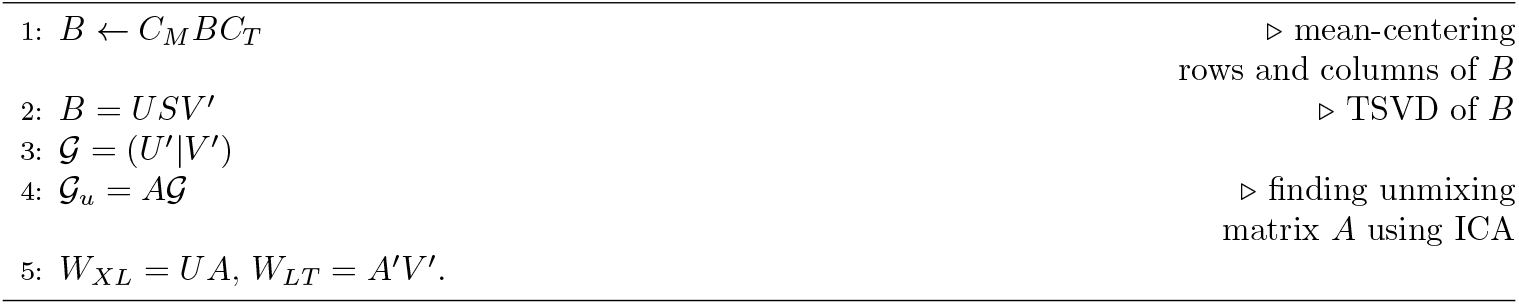

### 3.3 Contribution scores

We define the contribution scores for the SNPs and traits, respectively, as

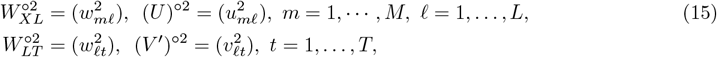

where we use the same notation for the weight matrices as before and the superscripted denotes the Hadamard (entrywise) exponent. By construction, both *W*_*LT*_ and *V* ^′^ are semiorthogonal, in the sense of satisfying 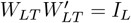 and *V* ^′^*V* = *I*_*L*_, where *I*_*L*_ is the identity matrix of size *L*, but 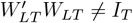 and *V V* ^′^ ≠ *I*_*T*_ (and analogously for *W*_*XL*_ and *U*). Therefore, if the total weight were uniformly distributed across *W*_*LT*_ and *V* ^′^, then every entry of 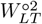 and (*V* ^′^) ^° 2^ would be given by 1*/T*. Thus, one way to characterize a latent is to record all traits for a given latent such that *w*_*𝓁t*_, *v*_*𝓁t*_ > 1*/T*. This is the default criterion we use for the GUIDE network visualization module in our software package. Taking the union of all traits with greater than uniform weights can also indicate the fraction of traits that are significantly represented by the latent factors, as we have done in Section 1.2.

### 3.4 Genetic variance components

Given the truncated singular value decomposition (TSVD) 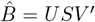, the unmixing matrix defined in Algorithm 1, *A*, we define the TSVD factor scores,

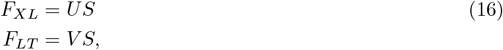

and the GUIDE factor scores,

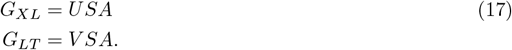

The corresponding definition for the GUIDE factor scores can be justified in several ways. First, the unmixing (rotation) matrix can only be applied after the scaling due to *S* is applied, since otherwise (if we had defined it as *UAS*, for example) *A* would have likely permuted at least some of the factors, so the singular values would have been applied to the wrong factors. Alternatively, we replace each of the matrices in Eq. (16) with their GUIDE counterparts: *U* ↦ *W*_*XL*_ = *UA, V* ↦ *W*_*LT*_ = *V A*, and *S* ↦ *A*^′^*SA*. Thus, *G*_*XL*_ = *W*_*XL*_*T* = *UAA*^′^*SA* = *USA* and *G*_*LT*_ = *W*_*LT*_ *T* = *V AA*^′^*SA* = *V SA*.

Writing *F*_*XL*_ = (*f*_*x𝓁*_), *F*_*LT*_ = (*f*_*𝓁t*_), *G*_*XL*_ = (*g*_*x𝓁*_), *G*_*LT*_ = (*g*_*𝓁t*_), the genetic variance components for each factor are then defined as,

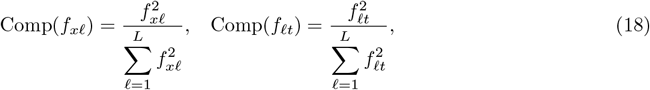

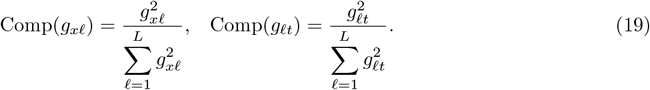

These scores are referred to as ‘cosine squared scores’ in, *e*.*g*., [6].

### 3.5 Weight strength significance of GUIDE models

We seek to estimate a “weight significance” or “weight strength value” (in short, a *w*-value) for any given 𝒳 → ℒ or ℒ → 𝒯 model weight by comparing it against the distribution of corresponding weights from all other possible models of the same data; in other words, by comparing the given weight with the corresponding matrix entries from all possible decompositions of the given summary statistics matrix (1). As mentioned in the main text, the *w*-value is the probability of randomly drawing the given weight value or a more extreme one given the distribution of corresponding weights from all possible models. Notably, though similar in spirit, *w*-values are not *p*-values because there is no specific null hypothesis weight against which a given model’s weight is compared, but rather the infinite set of other possible latent models of dimension *L*.

Up to scaling and signed permutations, all possible pairs of such matrices solving the decomposition problem (1), which includes the GUIDE solution as one of the possible decompositions, differ from each other by *L*-dimensional rotations (see the discussion in Sec. 3.1). (An animation showing examples of different models using simulated data can be found at https://guide-analysis.hail.is/.) Therefore, using a procedure similar to the one in [66] (though in that work the authors used different permutations as opposed to rotations), we used rotation matrices *R* ∈ O(*L*, ℝ) to sample from alternative models and thus obtained a distribution of alternative model weights for every 𝒳 → ℒ or ℒ → 𝒯 weight, from which *w*-values were computed. Since such *w*-values tended to be quite small and sampling enough models to measure them empirically would thus be computationally prohibitive, one solution is to sample 100-1000 models and use a Gaussian distribution with the same mean and standard deviation to compute the *w*-values.

To address the computational cost and errors associated with having to sample many models, we look for a closed form null distribution of the variance components. We first recognize that the *L*-vector of genetic variance components (see (18)) comprises a unit vector, and alternative models’ variance components are rotations of these, given by

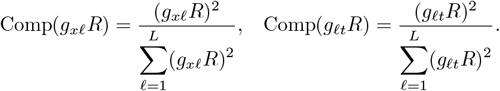

Sampling these is equivalent to sampling points uniformly from the unit (*L* − 1)-sphere. Writing *Z*_*𝓁*_ := *g*_*x𝓁*_*R*, we use a well known method for sampling points uniformly from a hypersphere [67]. Sampling *Z*_*𝓁*_ ∼ 𝒩 (0, 1), we then have that 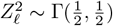 and 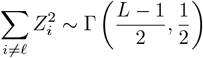, where Γ(*a, λ*) is the Gamma distribution with shape parameter *a* and rate parameter *λ*. We use the fact that if *X* ∼ Γ(*α, λ*) and *Y* ∼ Γ(*β, λ*), then 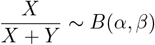, the beta distribution with shape parameters *α* and *β*. Therefore,

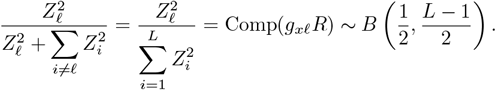

The same holds for the ℒ → 𝒯 variance components, Comp(*g*_*𝓁t*_*R*). Therefore, sampling variance components from this beta distribution provides a closed-form method for sampling alternative models’ variance components to compute the GUIDE model’s *w*-values efficiently and precisely. See Fig. S16, which computationally verifies that the mean and standard deviations of the *p*-values computed using the sampling method converge to the closed-form values as the number of samples increases. Fig. S17 shows probability-probability (P-P) plots of observed GUIDE 𝒳 → ℒ and ℒ → 𝒯 values versus the − log_10_(*w*) values drawn from the theoretical null distribution described above.

Referring to Sec. 3.3, we note that 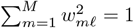 and 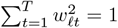, that is, the *M* -vector of 𝒳 → ℒ contribution scores and *T* -vector of ℒ → 𝒯 contribution scores are unit vectors. Thus, analogously with the above arguments, sampling contribution scores from alternative models is equivalent to sampling points from the unit (*M* − 1)- or (*T* −1)-sphere, which could therefore be sampled from the beta distributions, 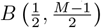 and 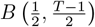, respectively.

As noted in Discussion, the effects of linkage disequilibrium (LD) tend to inflate model weights, which informed our prescription to LD prune the summary statistics before inputting into GUIDE. We analyzed this behavior in our simulations, where we showed that data with high LD tended to inflate estimated *w*-values, with characteristic ‘plateauing’ behavior seen in the P-P plots, wherein variants in LD loaded onto the same latent factor with approximately equal, and inflated, *w*-values. Simulations with low LD (corresponding to LD-pruned data) showed more limited inflation and plateauing and qualitatively agreed with the results observed in the real data in Fig. S17. Fig. S18 shows the P-P plots of the simulation’s ground truth *w*-values, as well as the GUIDE and TSVD models’ plots with LD and with LD-pruned simulated data. While both methods exhibited the above inflationary and plateauing behavior, the effects were consistently smaller for GUIDE.

We note that this method for computing *w*-values could be used for any latent representation model and is independent of the GUIDE model. To be clear, the method described above estimates *w*-values for a particular latent representation of the given 𝒳 → 𝒯 data (the summary statistics) relative to all possible latent representations of dimension *L*, as opposed to estimating those for the given latent representation relative to all possible summary statistics matrices. This is in line with our intended application for these *w*-values, which is assessing the weight strength significance of a given GUIDE model weight relative to a distribution of corresponding weights from all possible competing models. See Sec. S1 for an analysis of the the effects of standard errors on the GUIDE model weights.

The *w*-values of set of nodes in a given layer loading onto a node in an adjacent layer (*e*.*g*., a set of variants on a particular latent factor, or a set of latent factors on a particular trait) (Fig. 1) can be combined using the harmonic mean (*cf*. [68]). For example, for a set of *L*_1_ latent factors loading onto a given trait *t*, the aggregated or harmonic mean *w*-value,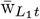, is given by,

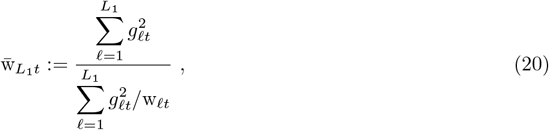

where w_*𝓁t*_ is the *w*-value for the weight between latent *𝓁* and trait *t*.

### 3.6 Genetic analysis of traits using GUIDE factors

Given a GUIDE model and a trait of interest, the top contributing latent factors are first determined based on their genetic variance components. Each latent factor is then characterized and interpreted by rank ordering the SNPs and traits that load onto it with the highest contribution scores, which can be visualized using stacked plots, as in Fig. 5, for example.

For further analysis, one may next ‘forward propagate’ from the given latent factor to the top contributing trait and then ‘back propagate’ to that trait’s top loading latent factor to compare the effects of these two latent factors. Alternatively, starting again from the first latent factor, one may back propagate to the top loading SNP (or group of SNPs) and then forward propagate to its top loading latent factor for a similar analysis. One may alternatively start with a given SNP or set of SNPs instead of a trait and proceed as above.

To isolate mechanisms that do not involve some other trait or process—*e*.*g*., as was done in Sec. 1.4 with FEV1 and smoking behavior—one may rank order, based on the genetic variance components, the latent factors for both traits and subtract the latter from the former. In Sec. 1.4 this was used to identify latent factors contributing to FEV1 that do not involve influencing risk for smoking behavior or addictive behavior.

To automate these analyses and improve interpretability of the results, we have added a function that uses the variance components, contribution scores, or the *w*-values described above with user-specified thresholds. First, it provides a list of traits which have at least one significant ℒ → 𝒯 weight, as determined by the given weight or *w*-value threshold. This gives a subset of traits for which the given GUIDE model could be said to have performed well.

It also provides the significant latent factors for each of those traits and the significant variants for each of those latent factors, each based on its own threshold. Additionally, a list of traits for each latent factor, with weights meeting the specific cutoff, is also given. In all, these allow for a more streamlined interpretation of the mechanism or pathway associated with each latent factor, and for the determination of the appropriate sets of variants associated with a trait of interest through the intermediate latent factors.

To aid in the visualization of the network structure resulting from these analyses we have also added a function that plots the network or a given portion of it, and which can also provide an animation of the same network over several different threshold values.

### 3.7 Stability of GUIDE latent factors

To check for consistency of the latent factors we divided in half the set of SNPs in the original UKB dataset, random assigning each SNP to one of two partitions, and ran GUIDE on each partition after bootstrapping to ensure our stability measures do not depend on the random partition. We compared each partition’s GUIDE model with GUIDE model on the full dataset, which we considered the ground truth. We rank ordered the genetic variance components for every trait to account for the permutation of the latent factor labels between every run of the model (the sign is not a problem in this case since genetic variance components are computed by squaring the weights, and so are positive by definition). To quantify the error in the genetic variance components (and by extension the network weights) we used the weighted absolute percent error (WAPE), given by,

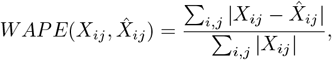

where *X*_*ij*_ are the ground truth values and 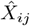 are those for the partitioned data.

For the randomly partitioned data the mean WAPE was around 8.5%, but seemed to have been driven up by a few outliers. As an example, for Alzheimer’s disease (AD), the WAPE was 2.0%, while for diabetes it was 1.3%.

We also partitioned the data based on even versus odd chromosome number, finding that the error, while on average higher for this nonrandom partition, was still fairly low for polygenic traits, but was higher for traits with top latent factors loading strongly on particular loci. For example, for AD, the WAPE for the odd chromosome partition (which includes the *APOE* gene on chromosome 19) was 6.5%, while for the even chromosome partition it was 24.0%. In contrast, for diabetes these were 11.7% and 10.6%, respectively.

We also randomly partitioned the traits and repeated the above analysis, finding that partitioning over traits resulted in higher errors (mean of 27%). This is because the latent factors are much more sensitive to loss of information on the traits’ side, given that these tend to be much fewer than the SNPs (order of 1000 versus 10^6^). We verified this trend in our simulations, where we found an average WAPE of 5% for the SNP partitions and 25% for the trait partitions.

We also assessed the stability of GUIDE latent factors using *T* jackknife subsamples of the traits (*M ×* (*T −* 1) betas matrices for every subsample), building a GUIDE model for each, and computing a WAPE for every one of the *M* variance component *L*-vectors, using the original model’s corresponding variance component vector as the ground truth. The average WAPE across all variants was 4.71%. Similarly, we conducted the corresponding analysis for the variants using ⌈*M/*100 ⌉ block jackknife subsamples of the variants (using blocks of 100 variants, giving (*M −*100) × *T* betas matrices for every subsample), and computing a WAPE for every one of the *T* variance component *L*-vectors. The average WAPE across all traits was 4.17%.

A full list of WAPE values for all the above analyses together with the code used to generate them can be found at https://github.com/daniel-lazarev/GUIDE.

### 3.8 Measuring SNP set overlap

In order to compare the percent overlap of two SNP sets, *S*_1_ and *S*_2_, we used the Jaccard index, defined as,

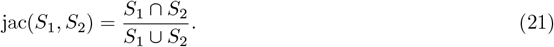

This was used in constructing Fig. S14.

### 3.9 Model selection with GUIDE

For a mixture of signals that includes non-Gaussian components as well as Gaussian ones, ICA will find a rotation to reconstruct the non-Gaussian components, but will include some random rotation for the Gaussian ones [9, 10]. Given this, if we have a dataset described by *L* latent factors corrupted by some Gaussian noise, and in our initial run of GUIDE we pick some *K* > *L*, the resulting unmixing matrix will include some components needed to rotate the *L*-dimensional signal (and noise), and some random rotation for the remaining *K* − *L* components (which only have noise). Now if we run ICA a second time, we get a different unmixing matrix. If we look at the cross-correlation of the two unmixing matrices, it will be a signed permutation for the parts that actually reconstruct the signal, but closer to 0 for the random rotation for the *K* − *L* “extra” components we added. This works because FastICA, the implementation of ICA we’re using, uses a random matrix to initialize the algorithm for finding the unmixing matrix [9, 10]. This gives a way to estimate the proper value of *L* to use in our model. To increase accuracy, we include an option to run ICA a third time and compare the resulting unmixing matrices in the same way.

### 3.10 Entropy analysis of GUIDE models

As an additional heuristic to guide model selection and assess the performance of the given model, we compute the entropies of the squared 𝒳 → ℒ or ℒ → 𝒯 weights, or contribution scores, defined by

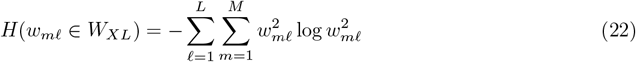

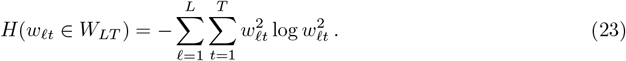

Computing entropies for every model with *L* latent factors traces out curves, such as in Fig. 4.

In general, entropy is maximized by the uniform distribution (signifying minimal information about the state of the system) and minimized by a probability distribution with all zeros except a one for one of the states (representing perfect knowledge of the state of the system) [21]. In the case of GUIDE weight matrices, higher entropies result from the signal being spread across more of the weights, while lower entropies result from sparser and more informative weight matrices.

For entropy plots such as those in Fig. 4, increasing *L* affects the entropy of a model in two competing ways. On one hand, it provides a larger space across which the signal could spread, increasing the entropy. On the other hand, it provides a richer space in which a successful latent representation would have enough ‘room’ to properly encapsulate the information in the data, which decreases the entropy. As *L* increases and approaches *T*, latent representation models could also have a tendency to maximize information trivially by simply reordering the traits, with the ℒ → 𝒯 weight matrices simply serving as permutation matrices.

## Code Availability

The functions for running GUIDE and analyzing its results, together with example analyses, are available at https://github.com/daniel-lazarev/GUIDE.

## Acknowledgements

The authors would like to thank Luke O’Connor, Raymond Walters, Mark J. Daly, and Nikolas Baya for helpful discussions and for their comments on the draft, and Yosuke Tanigawa and Manual A. Rivas for freely sharing their data. This work is supported by the Novo Nordisk Foundation (NNF21SA0072102), and US National Institutes of Health grants U01MH115727, R01MH101244, and R37MH107649. This research has been conducted using the UK Biobank Resource under Application Number 31063.

## Notes

https://github.com/daniel-lazarev/GUIDE

### Competing Interest Statement

Benjamin M. Neale is a member of the scientific advisory board at Deep Genomics and Neumora. The rest of the authors declare no conflicts of interest.

### Summary of Updates

- New method to estimate significance of learned model weights - New analyses of stability of GUIDE latent factors - New discussions of questions such as generalizability, reproducibility, and model significance, together with several new analyses surrounding these questions - Improved exposition and correction of typographical errors

