## Supplementary Information for "GUIDE deconstructs genetic architectures using association studies"

### 1 Supplementary Information

#### 1.1 Analysis of the errors in the GUIDE decomposition

Given a sum of statistically independent rank one matrices, can one recover the terms? How, and under what conditions? This decomposition is of the form

$$Z = \sum_{k=1}^{K} t_k x_k y_k' = XTY', \tag{1}$$

where $Z$ is an $M \times N$ data matrix (with $\text{rank}(Z) = K$), $X$ is an $M \times K$ matrix, $T$ is a $K \times K$ diagonal matrix, $Y$ is an $N \times K$ matrix, and $x_k$ and $y_k$ are the $M$- and $N$-dimensional column vectors of $X$ and $Y$, respectively. The latter three matrices are the estimands in our problem. Note that since $x_k$ and $y_k$ are independent high dimensional vectors, they are approximately orthogonal, but not exactly orthogonal due to sampling variation.

As an initial decomposition, we find the compact singular value decomposition (SVD), given by

$$Z = USV', \tag{2}$$

where $S$ is a $K \times K$ diagonal matrix consisting of the nonzero eigenvalues of $ZZ'$, $U$ is an $M \times K$ orthogonal matrix consisting of the eigenvectors of $ZZ'$ corresponding to the eigenvalues in $S$, and $V$ is an $N \times K$ orthogonal matrix consisting of the same for $Z'Z$.

Unlike the columns of $X$ and $Y$, those of $U$ and $V$ are guaranteed to be exactly orthogonal by the SVD. If $X$ and $Y$ were orthogonal, then $X = U$ and $Y = V$, and the SVD would give the correct decomposition into statistically independent components.

Nevertheless, since $XTY' = USV'$, we can write

$$U = X \underbrace{TY'VS^{-1}}_{\mathcal{M}_X} \tag{3}$$

$$V = Y \underbrace{TX'US^{-1}}_{\mathcal{M}_Y} \tag{4}$$

where $\mathcal{M}_X$ and $\mathcal{M}_Y$ are the true mixing matrices. These two equations define two ICA problems. After running ICA on $U$ and $V$, respectively, we obtain

$$U = \hat{X}\mathcal{M}_{\hat{X}} \tag{5}$$

$$V = \hat{Y}\mathcal{M}_{\hat{Y}} \tag{6}$$

where the hatted matrices denote the estimated quantities obtained for the corresponding estimands. Unlike the estimands, the estimators $\hat{X}$, $\hat{Y}$, $\mathcal{M}_{\hat{X}}$ and $\mathcal{M}_{\hat{Y}}$ have orthonormal columns by construction.

The purpose of our analysis in this section is to find the estimators for $X, T, Y$ in (1) so that $\hat{X}\hat{T}\hat{Y}' \approx XTY'$. Two of these, $\hat{X}$ and $\hat{Y}$, are given by the ICA in (5) and (6). How do we estimate $\hat{T}$? By the above we have $USV' = XTY' \approx \hat{X}\hat{T}\hat{Y}'$. By (5) and (6) we obtain

$$\begin{aligned} USV' &= \hat{X}\mathcal{M}_{\hat{X}}S\mathcal{M}_{\hat{Y}}'\hat{Y}' \approx \hat{X}\hat{T}\hat{Y}' \\ \hat{T} &\approx \mathcal{M}_{\hat{X}}S\mathcal{M}_{\hat{Y}}'. \end{aligned} \tag{7}$$

A small error term, one for each of $X$ and $Y$, and reflecting the effects of the sampling variation, allows us to relate each estimand to its estimator:

$$\begin{aligned} X &= \hat{X}(I_K + \mathcal{E}_X) \\ Y &= \hat{Y}(I_K + \mathcal{E}_Y) \end{aligned} \tag{8}$$

Using these relations and the fact that the estimated quantities $\hat{X}\hat{T}\hat{Y}' \approx XTY'$, we obtain a relation for $\hat{T}$ in terms of the errors and the true $T$:

$$\begin{aligned} XTY' &= \hat{X}(I_K + \mathcal{E}_X)T(I_K + \mathcal{E}_Y')\hat{Y}' \\ &= \hat{X}(T + \mathcal{E}_X T + T\mathcal{E}_Y' + \cancel{\mathcal{E}_X T \mathcal{E}_Y'})\hat{Y}' \approx \hat{X}\hat{T}\hat{Y}' \\ \Rightarrow \hat{T} &\approx T + \mathcal{E}_X T + T\mathcal{E}_Y'. \end{aligned} \tag{9}$$

Formula (7) provides a way to estimate $\hat{T}$, and (9) shows that its off-diagonal elements are only due to the sample variation since $T$ is diagonal. Note that while the off-diagonal elements of $\hat{T}$ are purely due to the error terms ($\mathcal{E}_X$ and $\mathcal{E}_Y$), its diagonal entries, while containing the true values of $T$, are also affected by the same.

We can also find the error term relating $\mathcal{M}_{\hat{X}}$ to $\mathcal{M}_X$ in terms of $\mathcal{E}_X$. Write

$$\begin{aligned} \mathcal{M}_X &= \mathcal{M}_{\hat{X}}(I_K + \mathcal{E}_{\mathcal{M}_X}) \\ \mathcal{M}_Y &= \mathcal{M}_{\hat{Y}}(I_K + \mathcal{E}_{\mathcal{M}_Y}). \end{aligned} \tag{10}$$

We will only explicitly write the calculations for $X$, as those for $Y$ are analogous. From (3), (8), and (10) we have

$$\begin{aligned} U = X\mathcal{M}_X &= \hat{X}(I_K + \mathcal{E}_X)\mathcal{M}_{\hat{X}}(I_K + \mathcal{E}_{\mathcal{M}_X}) \\ &= \hat{X}\mathcal{M}_{\hat{X}} + \hat{X}\mathcal{E}_X\mathcal{M}_{\hat{X}} + \hat{X}\mathcal{M}_{\hat{X}}\mathcal{E}_{\mathcal{M}_X} + \cancel{\hat{X}\mathcal{E}_X\mathcal{M}_{\hat{X}}\mathcal{E}_{\mathcal{M}_X}} \\ &\approx U + \hat{X}\mathcal{E}_X\mathcal{M}_{\hat{X}} + \hat{X}\mathcal{M}_{\hat{X}}\mathcal{E}_{\mathcal{M}_X} \end{aligned}$$

where we used (5) in the last step, and canceled the term that is second order in the error in the penultimate one. This gives an expression for the error term for the estimated mixing matrix in terms of the error for the estimated $X$:

$$\begin{aligned} \hat{X}\mathcal{M}_{\hat{X}}\mathcal{E}_{\mathcal{M}_X} &\approx -\hat{X}\mathcal{E}_X\mathcal{M}_{\hat{X}} \\ \mathcal{E}_{\mathcal{M}_X} &\approx -\mathcal{M}'_{\hat{X}}\mathcal{E}_X\mathcal{M}_{\hat{X}} \end{aligned} \tag{11}$$

Plugging (11) into (10) we obtain:

$$\begin{aligned} \mathcal{M}_X &\approx \mathcal{M}_{\hat{X}} - \mathcal{M}_{\hat{X}}\mathcal{M}'_{\hat{X}}\mathcal{E}_X\mathcal{M}_{\hat{X}} \\ \mathcal{M}_X &\approx (I_K - \mathcal{E}_X)\mathcal{M}_{\hat{X}} \end{aligned} \tag{12}$$

Using formulas (8) and (12) we can check that the approximations we made are justified by verifying that the two expressions for $U$ in (3) and (5) match up to a negligible error term:

$$\begin{aligned} X\mathcal{M}_X &\approx \hat{X}(I_K + \mathcal{E}_X)(I_K - \mathcal{E}_X)\mathcal{M}_{\hat{X}} \\ &= \hat{X}(I_K - \mathcal{E}_X^2)\mathcal{M}_{\hat{X}}, \end{aligned}$$

so using the above approximations does indeed reproduce the relation $X\mathcal{M}_X = \hat{X}\mathcal{M}_{\hat{X}}$ up to $O(\mathcal{E}_X^2)$.

#### 1.2 Kurtosis as a measure of performance

Table 1 and Fig. 1 compare the kurtosis for the latent factors produced by DeGAs versus that of GUIDE for the 'all' dataset. Kurtosis is a computationally favorable proxy measure for non-Gaussianity (or negentropy), which is the cost function maximized (or minimized) by ICA algorithms [1, 2]. As discussed in more depth in Section 1.4, this is one way of assessing the performance of the ICA step of our algorithm.

Table 1

| Kurtosis ('all' dataset) | DeGAs | GUIDE |
|---|---|---|
| Mean | 185.87 | 813.51 |
| Median | 116.63 | 685.93 |
| Range | 826.14 | 2108.86 |
| Std Dev | 184.04 | 635.65 |
| Max | 838.18 | 2126.20 |
| Min | 12.03 | 17.33 |

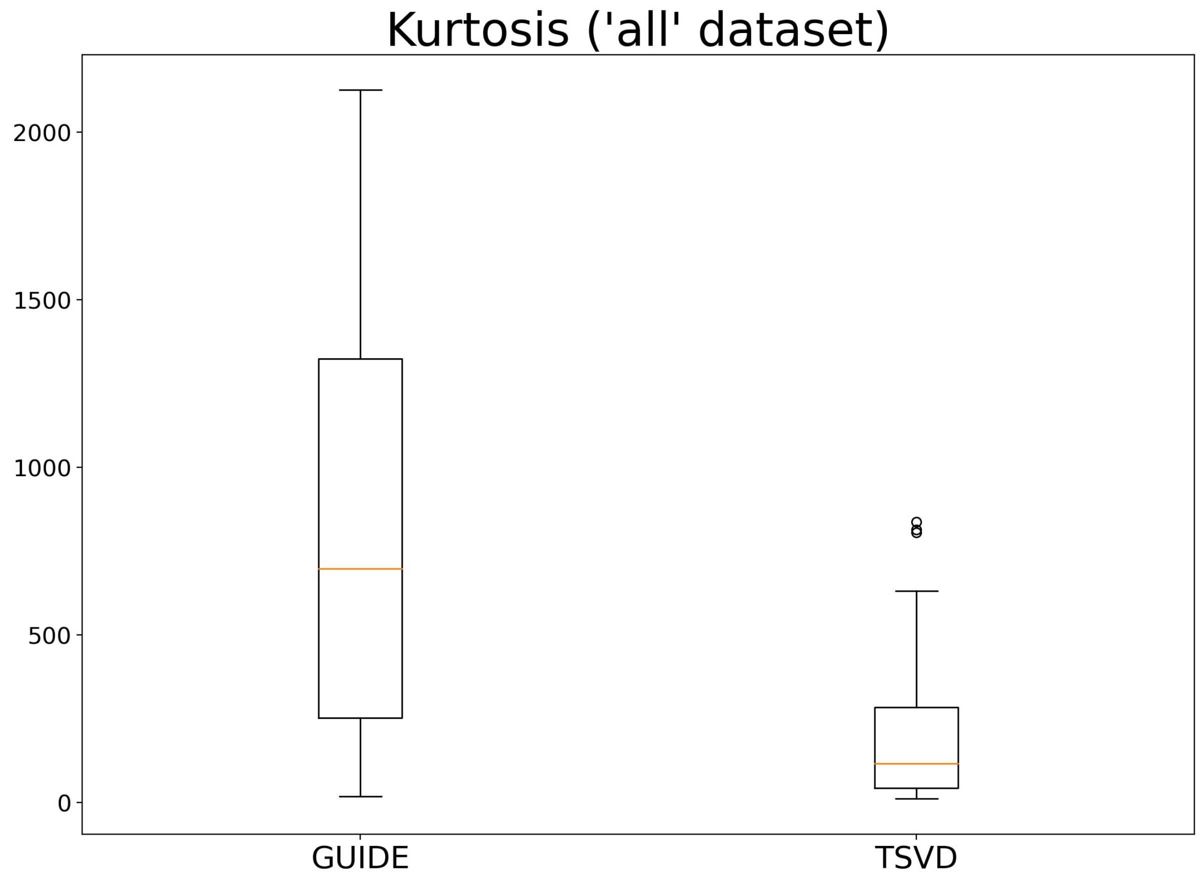


Figure 1: Center line, median; box limits, upper and lower quartiles; whiskers, 1.5x interquartile range; points, outliers.

#### 1.3 Independence of latent factors

As mentioned in the main text, ICA maximizes nongaussianity by maximizing negentropy, defined as the difference between the entropy of a Gaussian distribution (which is the maximum entropy among all distributions with equal variance) and the entropy of the given distribution. ICA can equally be interpreted as minimizing mutual information, given by

$$I(X,Y) = \sum_{x,y} p(x,y) \log \left( \frac{p(x,y)}{p(x)p(y)} \right),$$

where $p(x,y)$ is the joint distribution for the random variables $X,Y$ and $p(x), p(y)$ are the corresponding marginal distributions [2]. Intuitively, mutual information measures how far $p(x,y)$ is from being independent. Thus, in the context of GUIDE, ICA finds latents factors that mutually minimize mutual information, and are in this sense 'maximally independent'. Notably, this notion of independence is stronger than merely decorrelation of factors, since the former includes higher order (as opposed to only second-order) and nonlinear effects [3].

#### 1.4 Heritability and trait enrichment of GUIDE factors

We prepared a set of minimally pruned UK Biobank data with 2,692,064 million SNPs and 4,357 phenotypes (see Methods for details), which we will refer to as the "UKB dataset". We then built a GUIDE model for this dataset with $L = 200$ (see Section 3.9 in Methods for a discussion on how this number was estimated using GUIDE's model selection algorithm).

Table 2 and Fig. 2 compare the heritabilities of the latent factors against those of the traits for the UKB dataset. The latent factors carry approximately the same total heritability as the traits, and so on average have a much higher heritability per trait (Methods).

Table 2: Statistics for the (normalized) heritabilities of the final traits versus the latent factors.

| | Final trait $h^2$ | Latent factor $h^2$ |
|---|---|---|
| Mean | 0.0002295157 | 0.0057803467 |
| Median | $9.5080 \times 10^{-8}$ | $3.2220 \times 10^{-4}$ |
| Mode | $9.5080 \times 10^{-8}$ | $2.7972 \times 10^{-7}$ |
| Standard deviation | 0.007022 | 0.01813 |
| Maximum | 0.26688 | 0.13398 |
| Minimum | $9.3536 \times 10^{-8}$ | $2.7972 \times 10^{-7}$ |

Fig. 3 shows the genetic genetic variance component values for the UKB $Z > 4$ dataset, for which close to 90% of GUIDE weights are larger than the corresponding TSVD weights (for the other 10% they are comparable, likely reflecting that this dataset does not include the necessary SNPs to inform the latent factors for those specific traits). Moreover, 29.8% of GUIDE latent weights have genetic genetic variance component scores above 0.9, as compared to 3.5% of TSVD latents factors.

#### 1.5 Supplementary figures and tables

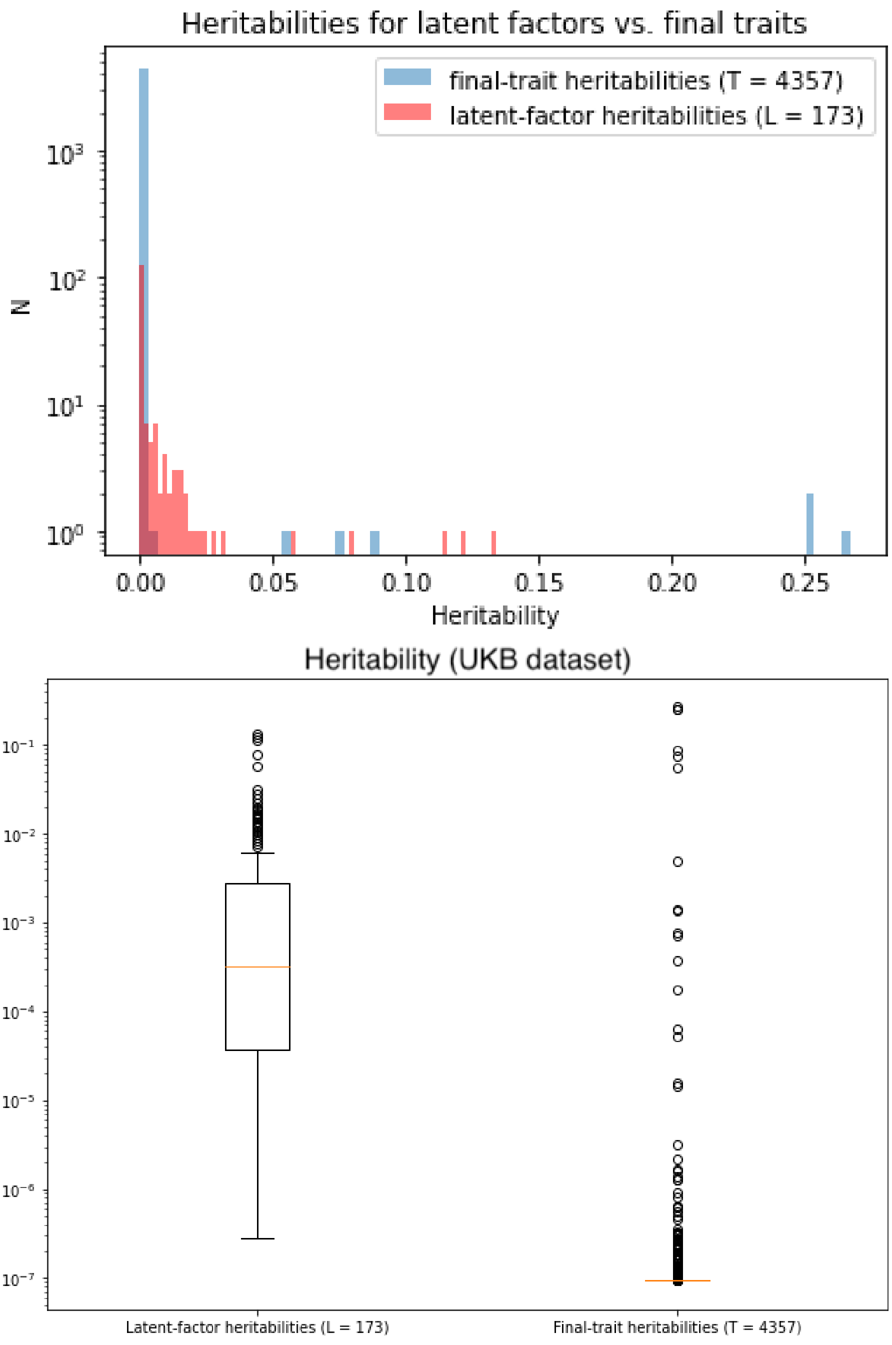


Figure 2: A comparison of the heritabilities for the final traits versus those for the latent factors, showing that latent factors capture more heritability on average, and moreover, despite being far fewer in number, capture the same total heritability for all traits. Center line, median; box limits, upper and lower quartiles; whiskers, 1.5x interquartile range; points, outliers.

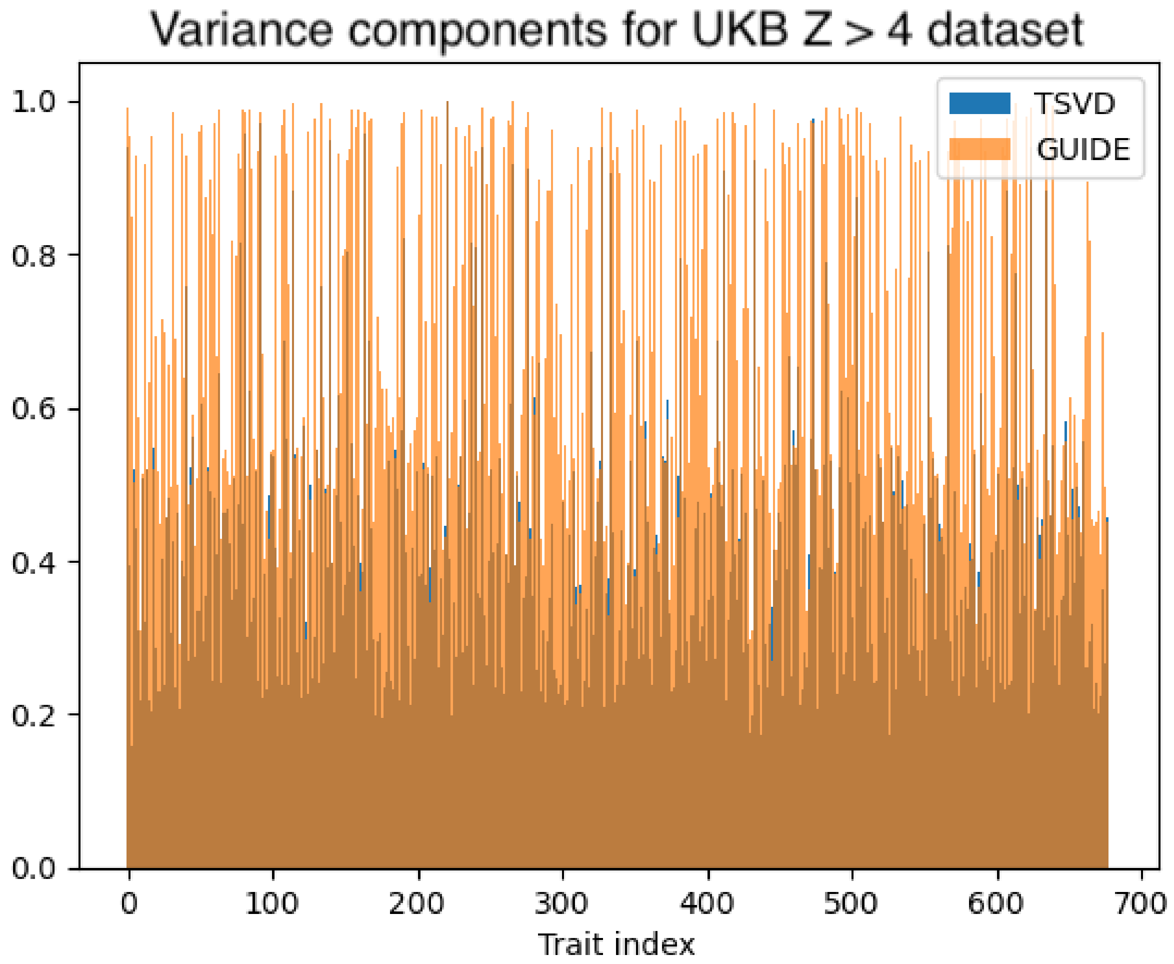


Figure 3: Here we plot, as in Fig. 3, the genetic variance components for traits in the 'all' dataset from [4] for the top three latent factors, but this time for all 2,138 traits. In this plot, the GUIDE latent factors were larger than their TSVD counterparts in 89.5% of cases.

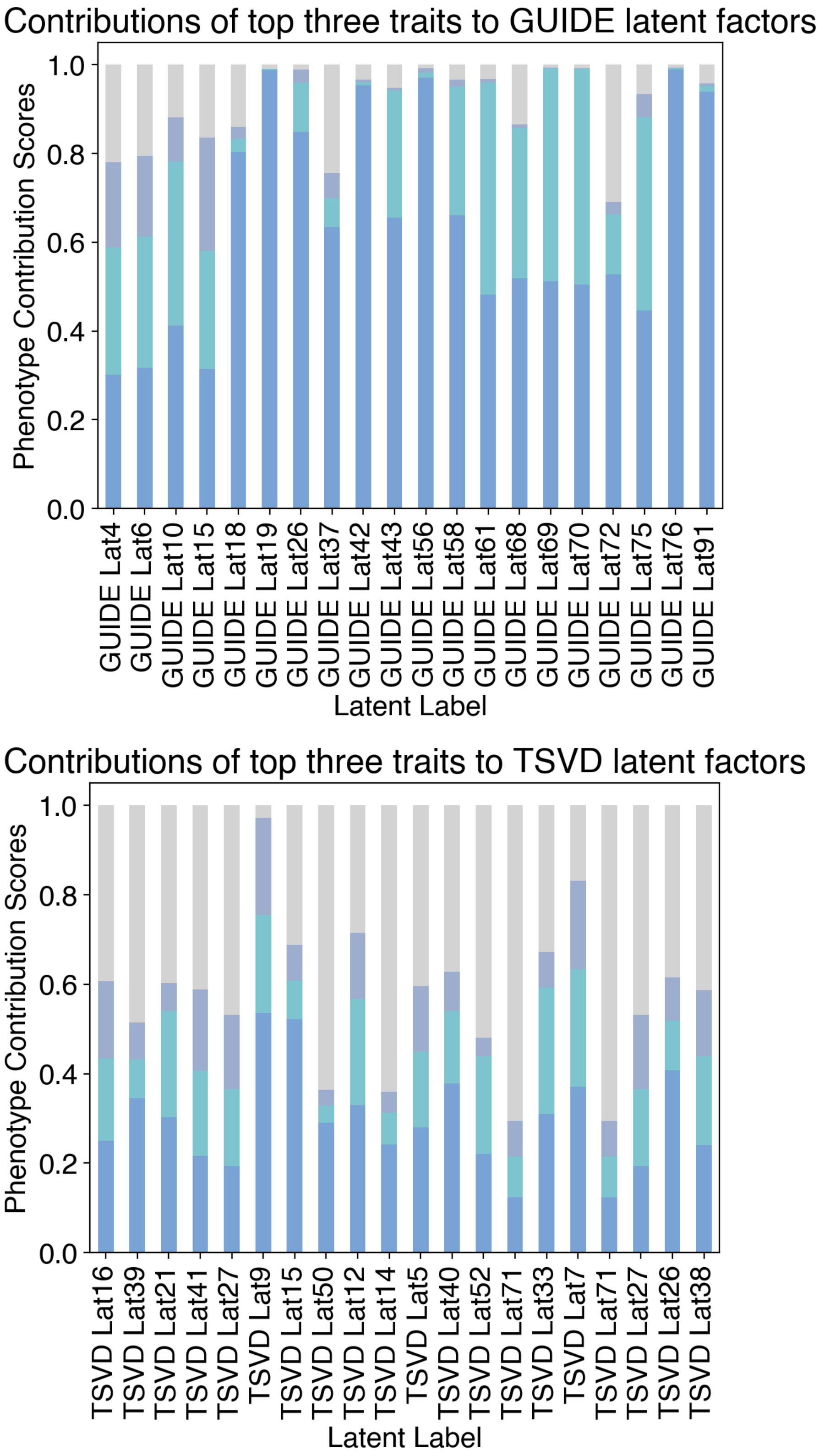


Figure 4: Stacked bar plots comparing, for each latent factor, the contributions from the top loading traits. See Fig. S8 for the variance explained curves for all 100 latent factors for GUIDE and for TSVD.

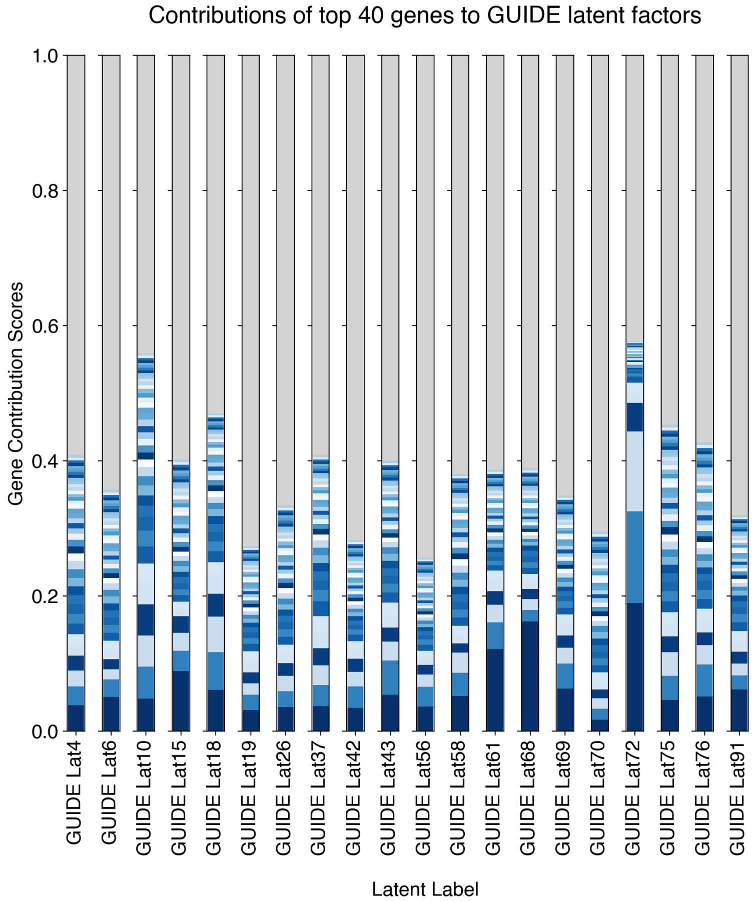


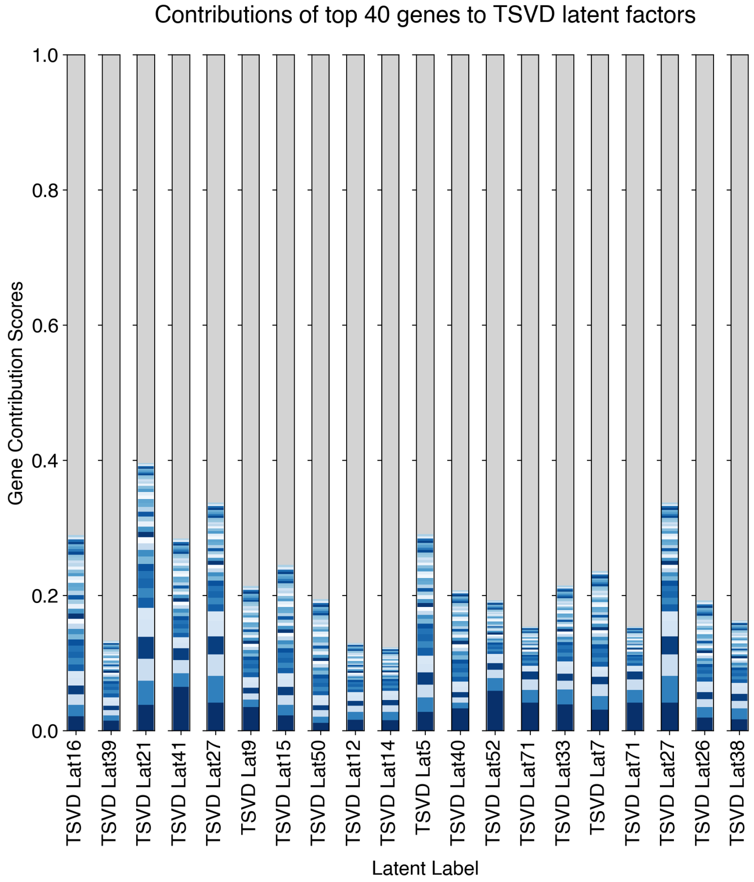


Figure 5: Stacked bar plots comparing, for each latent factor, the contributions from the top loading SNPs.

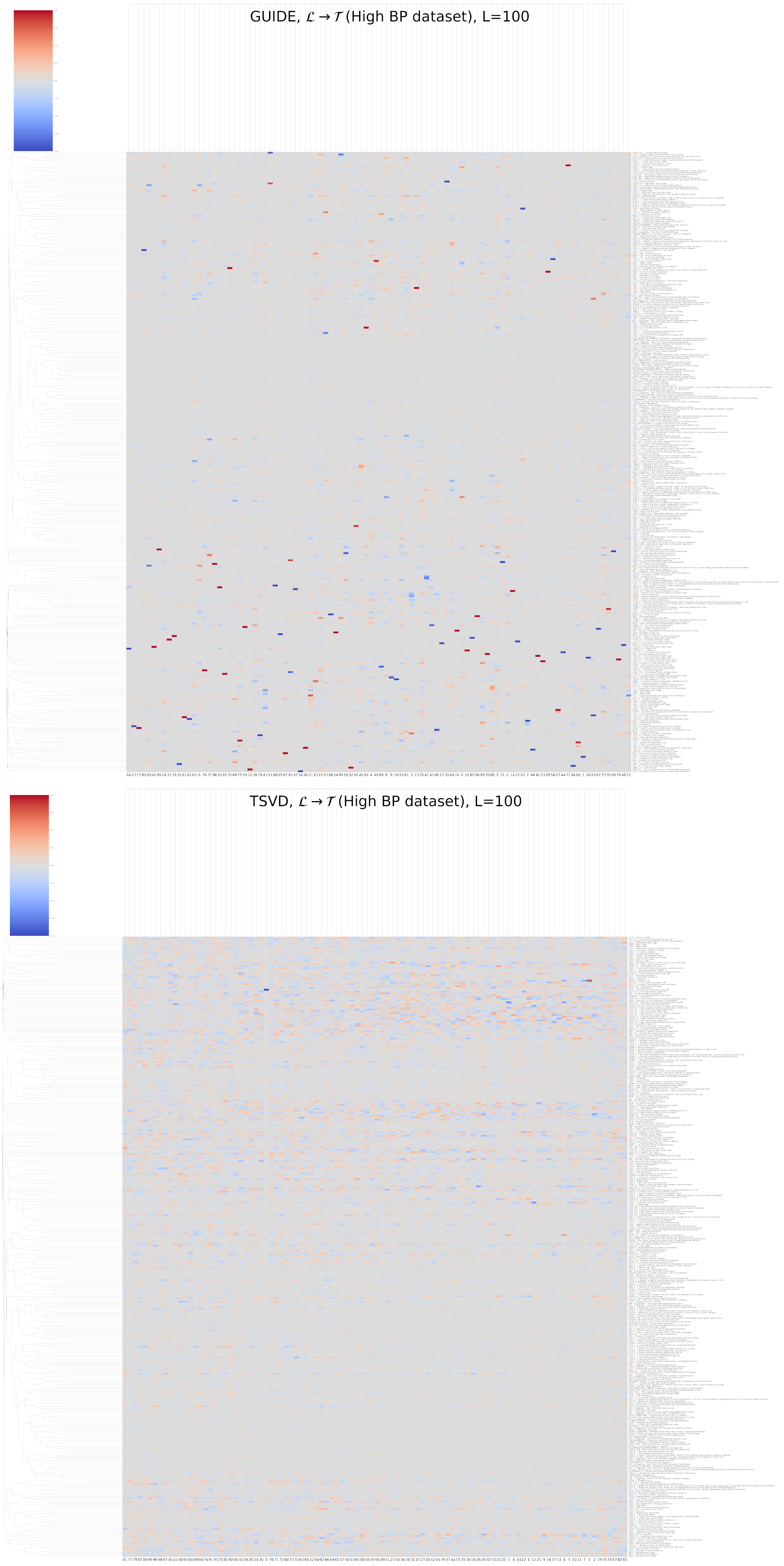


Figure 6: GUIDE $\mathcal{L} \to \mathcal{T}$ weights for loci significant for the "High Blood Pressure" dataset for for $L = 100$ compared with the corresponding TSVD $\mathcal{L} \to \mathcal{T}$ weights.

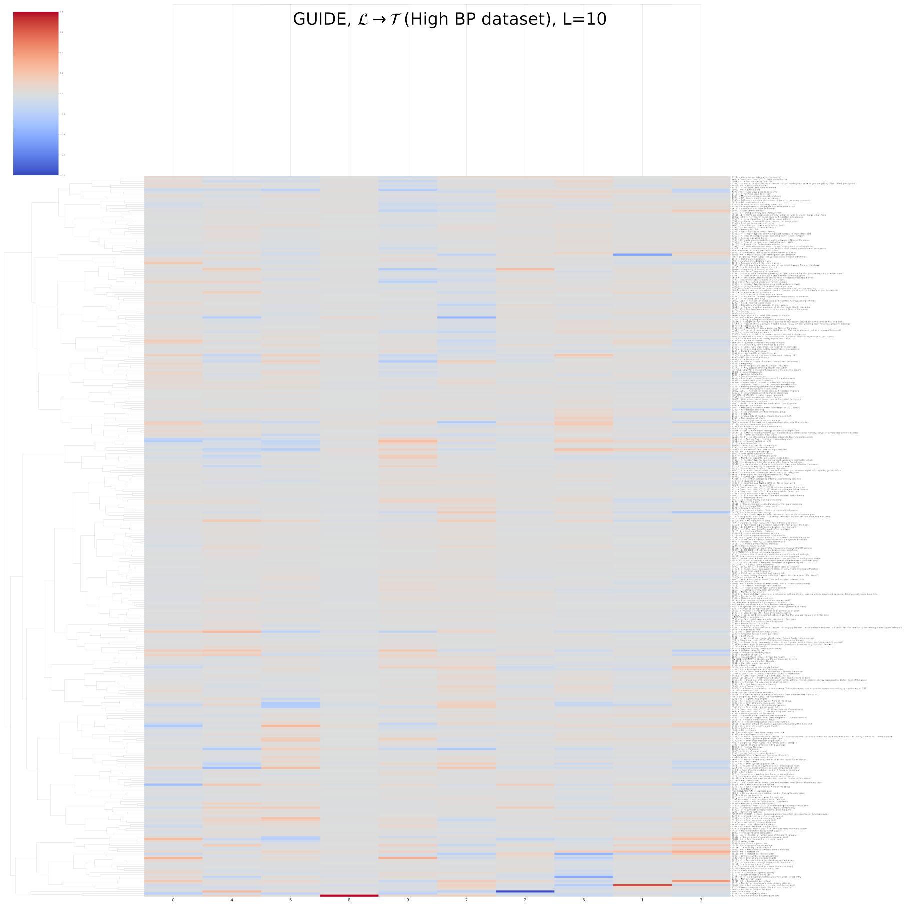


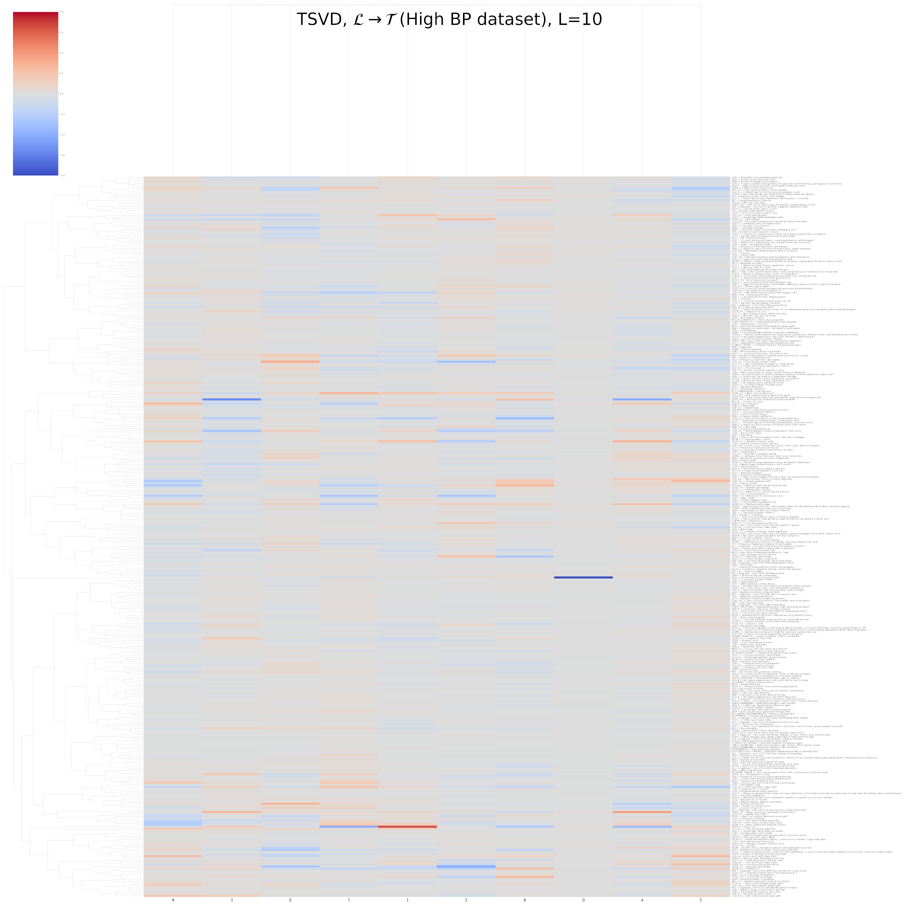


Figure 7: GUIDE $L \to T$ weights for loci significant for the "High Blood Pressure" dataset for for $L = 10$ compared with the corresponding TSVD $\mathcal{L} \to \mathcal{T}$ weights.

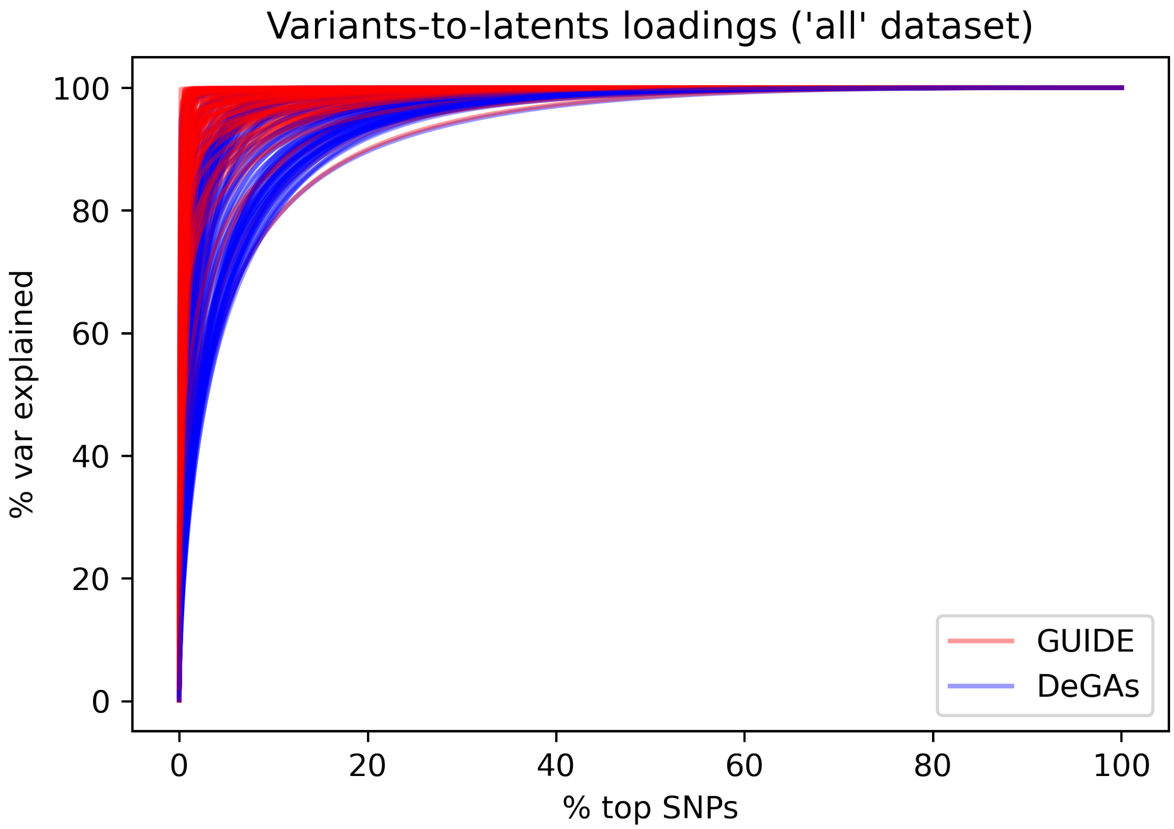


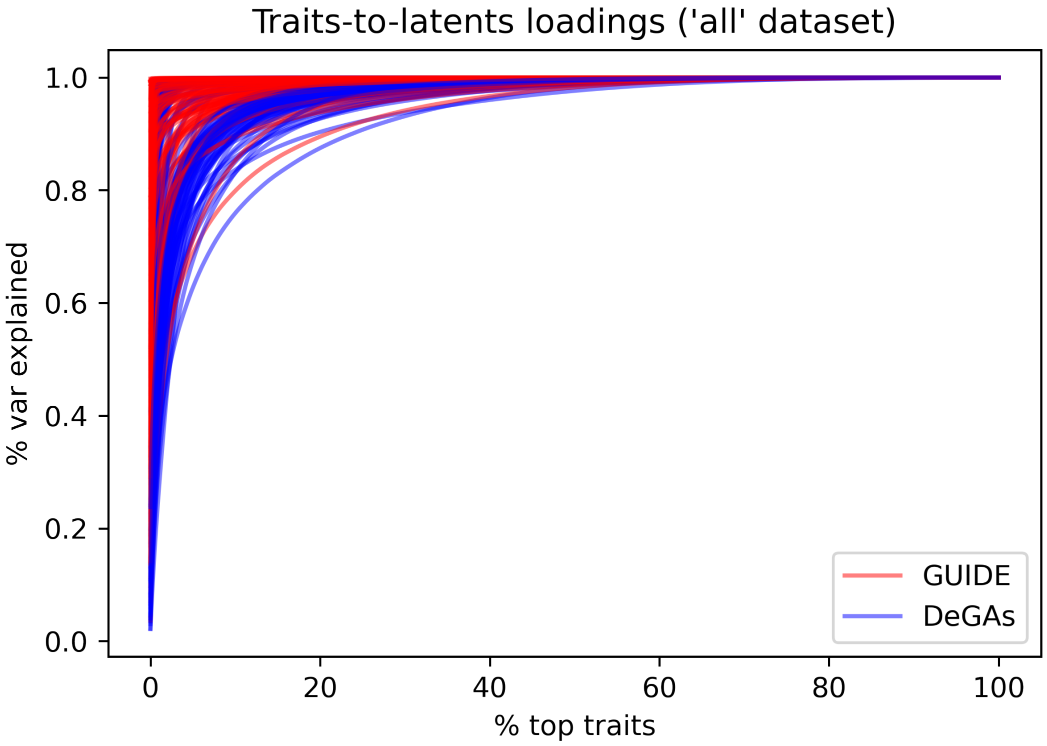


Figure 8: Variance explained plots comparing GUIDE with DeGAs as a function of the rank-ordered (based on $\mathcal{X} \rightarrow \mathcal{L}$ or $\mathcal{T} \rightarrow \mathcal{L}$ weights) SNPs or traits for every latent factor. Each line corresponds to a different latent factor. For GUIDE, most of the variance is explained by fewer SNPs and fewer traits for most of the latent factors. Results shown for the 'all' dataset [4] consisting of 235,907 variants and 2,138 traits, and both models were built using 100 latent factors.

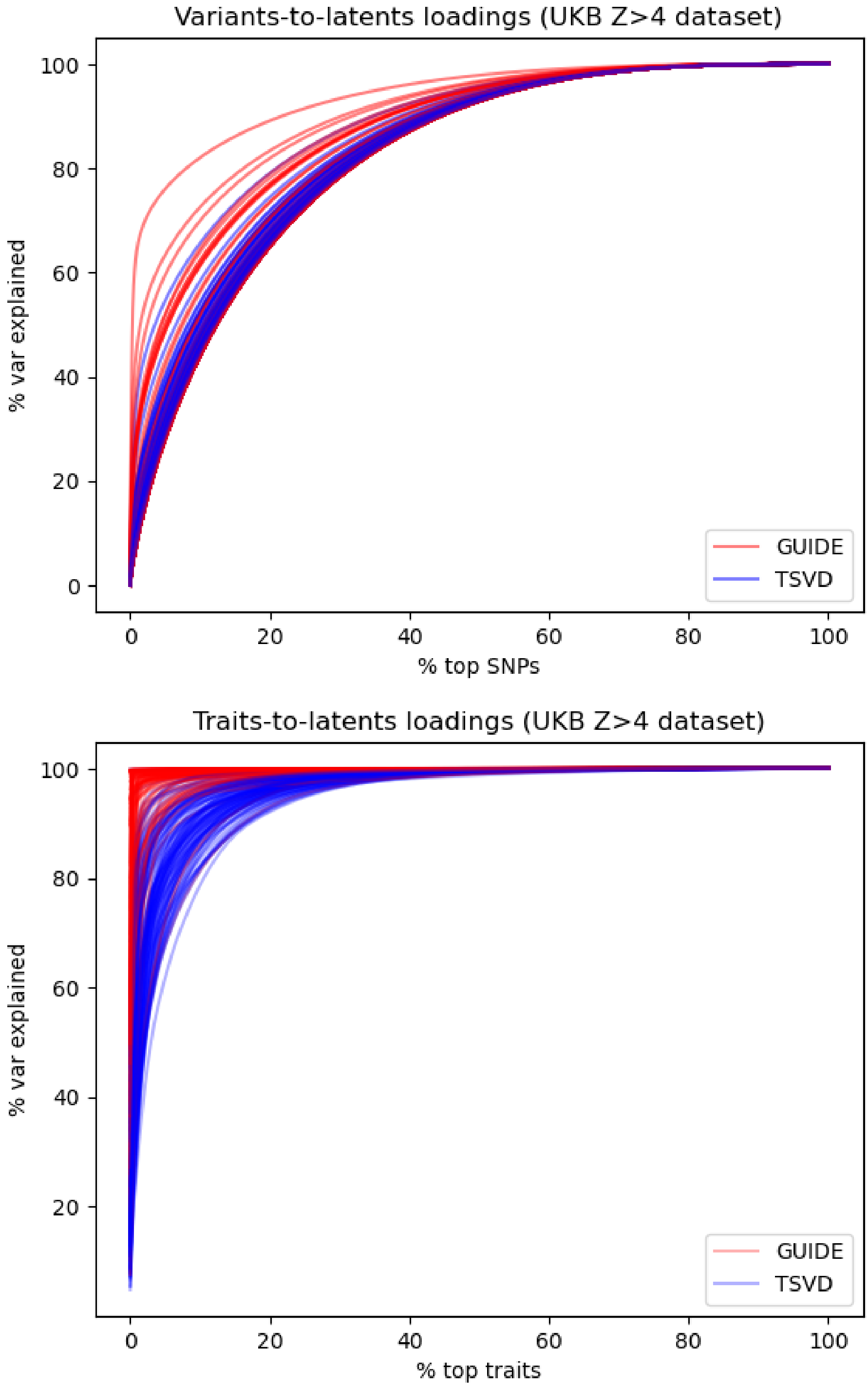
Variants-to-latents loadings (UKB Z>4 dataset)
% var explained
% top SNPs
GUIDE
TSVD
Traits-to-latents loadings (UKB Z>4 dataset)
% var explained
% top traits
GUIDE
TSVD

Figure 9

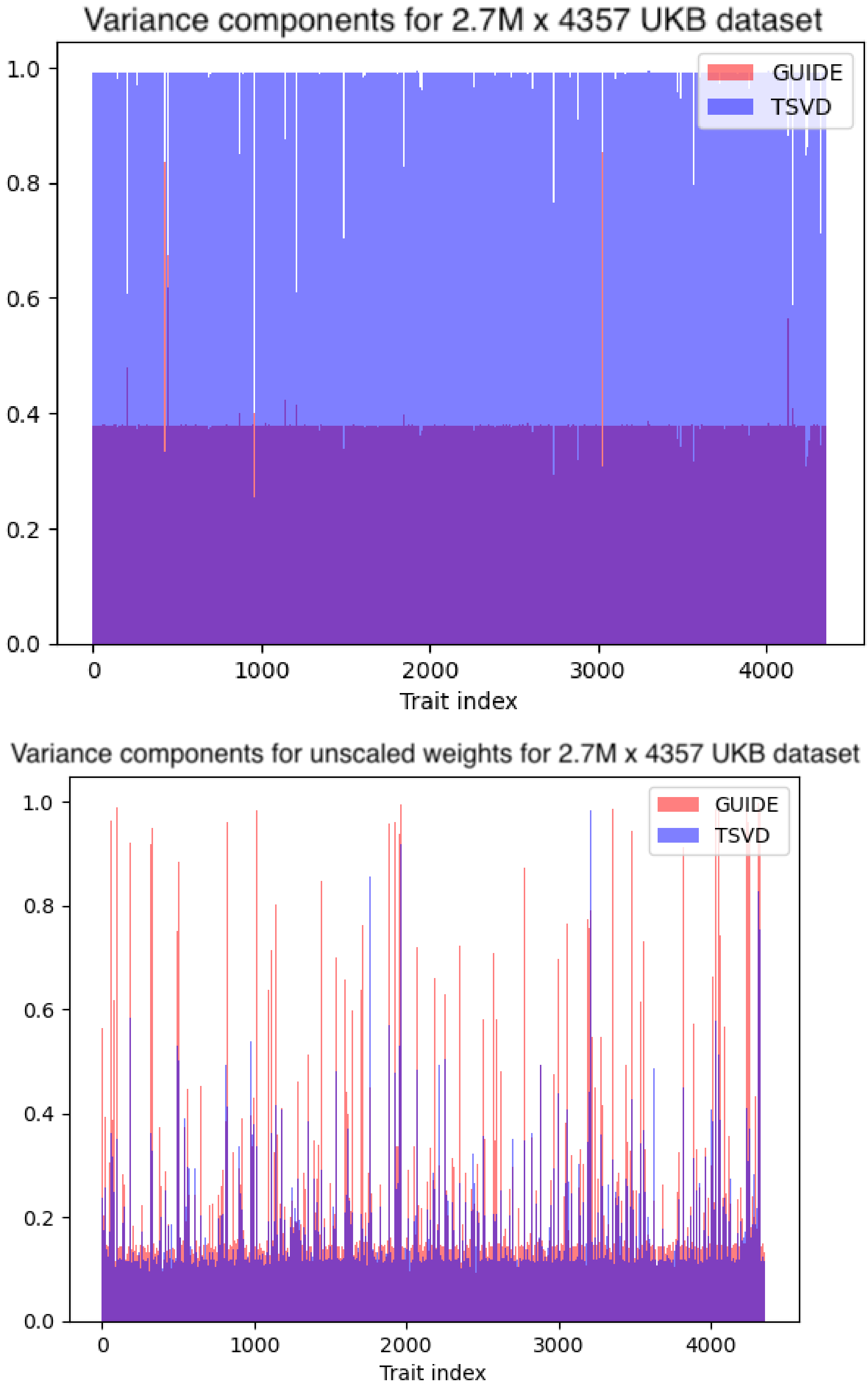


Figure 10: For the unpruned UK Biobank dataset, the TSVD latent factors have genetic variance components that are higher than their GUIDE counterparts in 98.9% of cases, unlike the results for $Z > 4$ dataset in Fig. 3. As discussed in the main text, this is likely due to LD and genetic covariance, which future work will hopefully address. Unsurprisingly, when we calculate the same plot but using the variance contribution score that is unweighted by the singular values (that is, $V$ instead of $VS$ and $VA$ instead of $VSA$ in Equations 16 and 17), the opposite is the case, with the GUIDE factors outweighting the TSVD ones 85.1% of the time. This is expected since the ICA basis was found for the unscaled data.

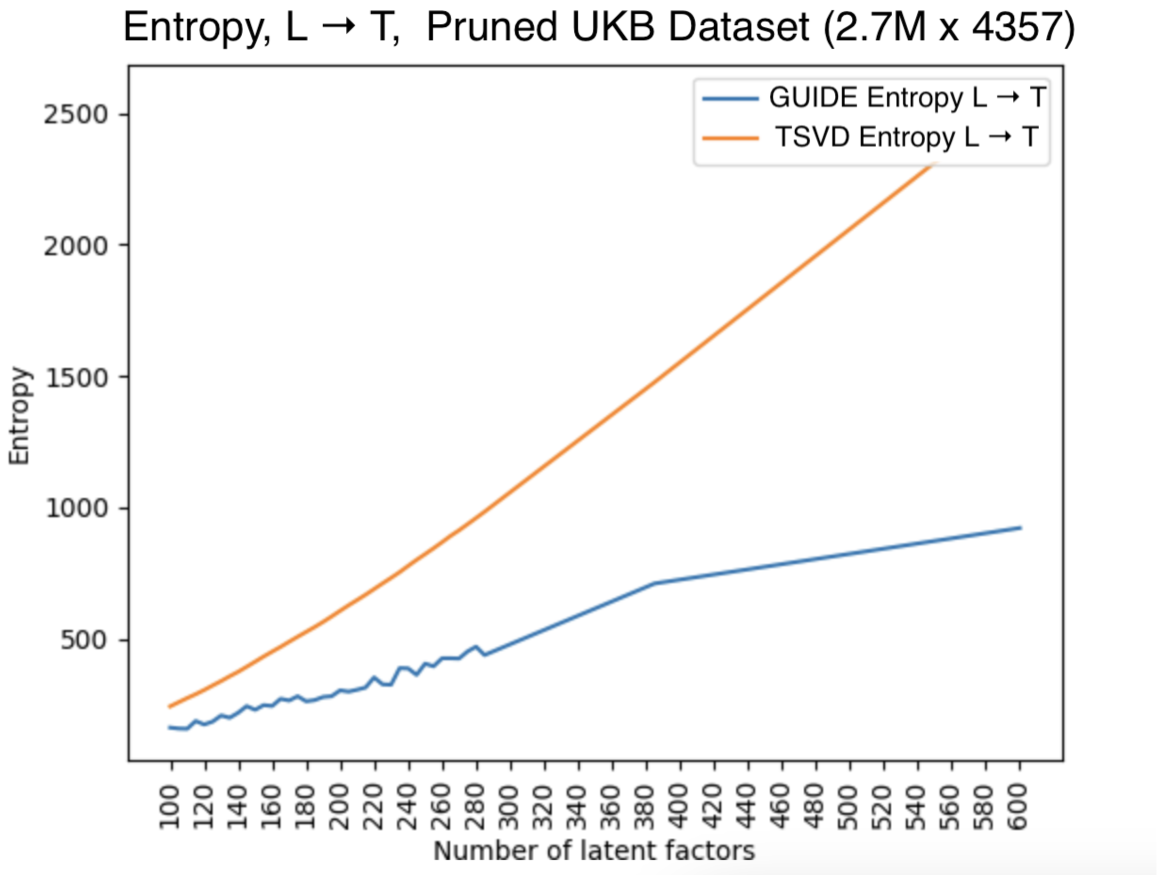


Figure 11: The entropy of the $\mathcal{L} \to \mathcal{T}$ weights as a function of the number of latent factors in the each model for the full UKB dataset (2,692,064 million SNPs and 4,357 phenotypes).

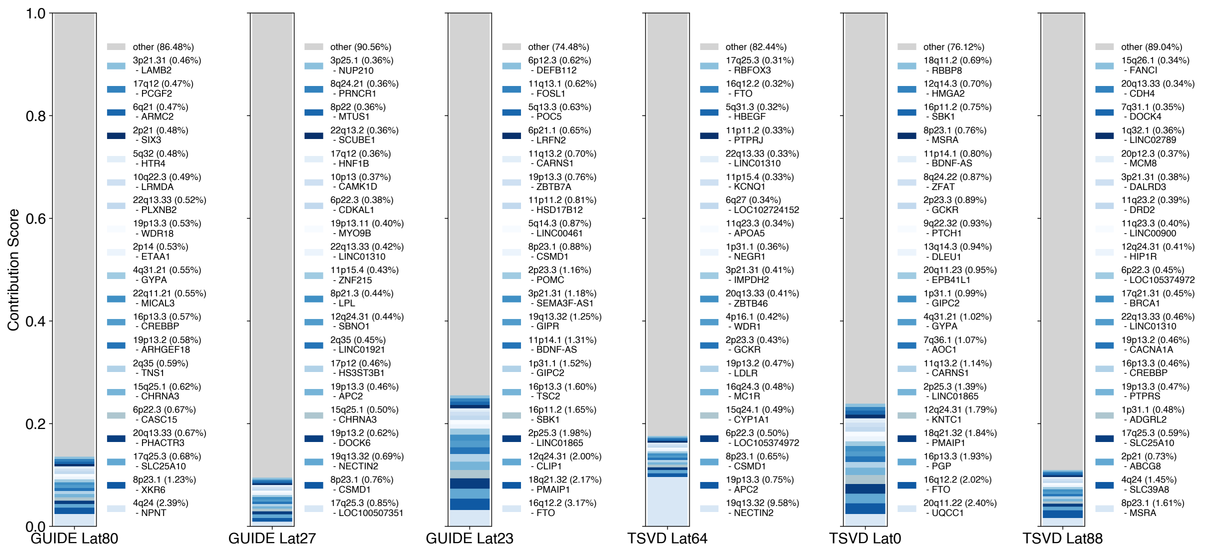


Figure 12: Top loci implicated by the leading GUIDE and TSVD latent factors for the traits ‘Number of cigarettes smoked daily’ and ‘Number of cigarettes previously smoked daily’.

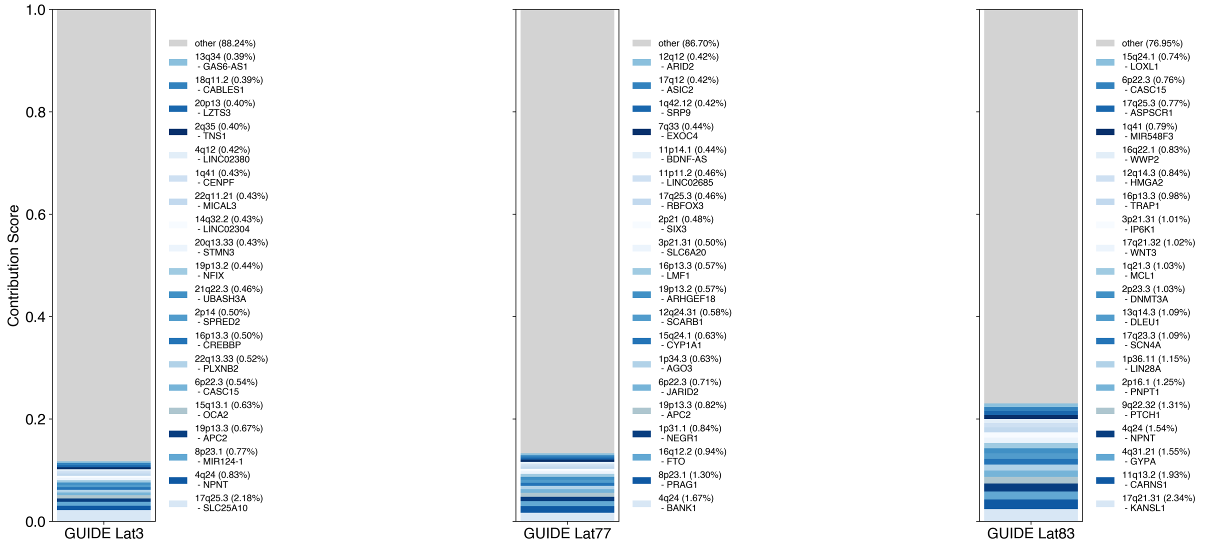


Figure 13: GUIDE top latent factors for the 'all' dataset without the smoking-related traits. GUIDE latent factors for the with- and without-smoking models were matched using the top loading traits for each factor, which was consistent between the top latent factors for each model.

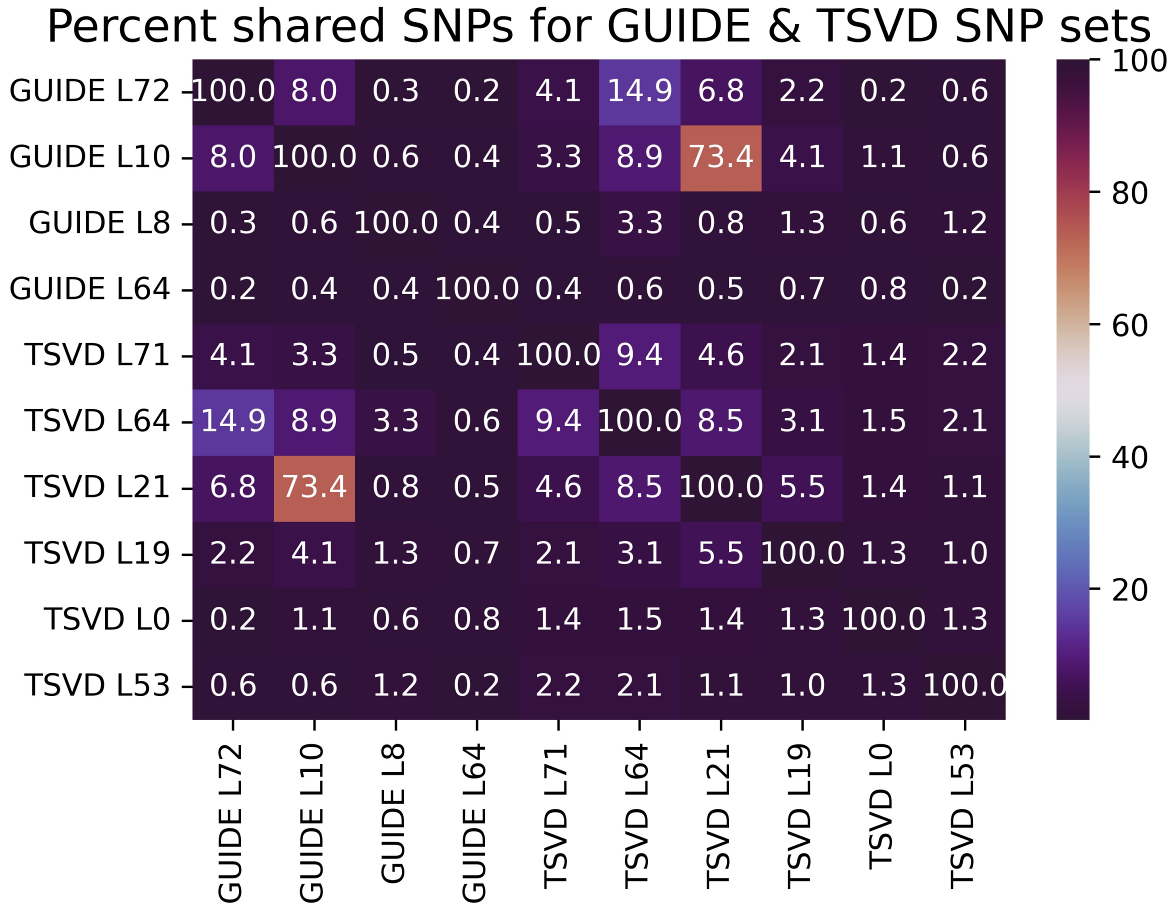


Figure 14: Percent common SNPs (as measured by the Jaccard index, Methods) for the top 500 SNPs corresponding to the GUIDE and TSVD latents in the subplots of Figure 3
. While the general cholesterol factors (GUIDE latent 10 and TSVD latent 21) share a many SNPs, all other factors do not.

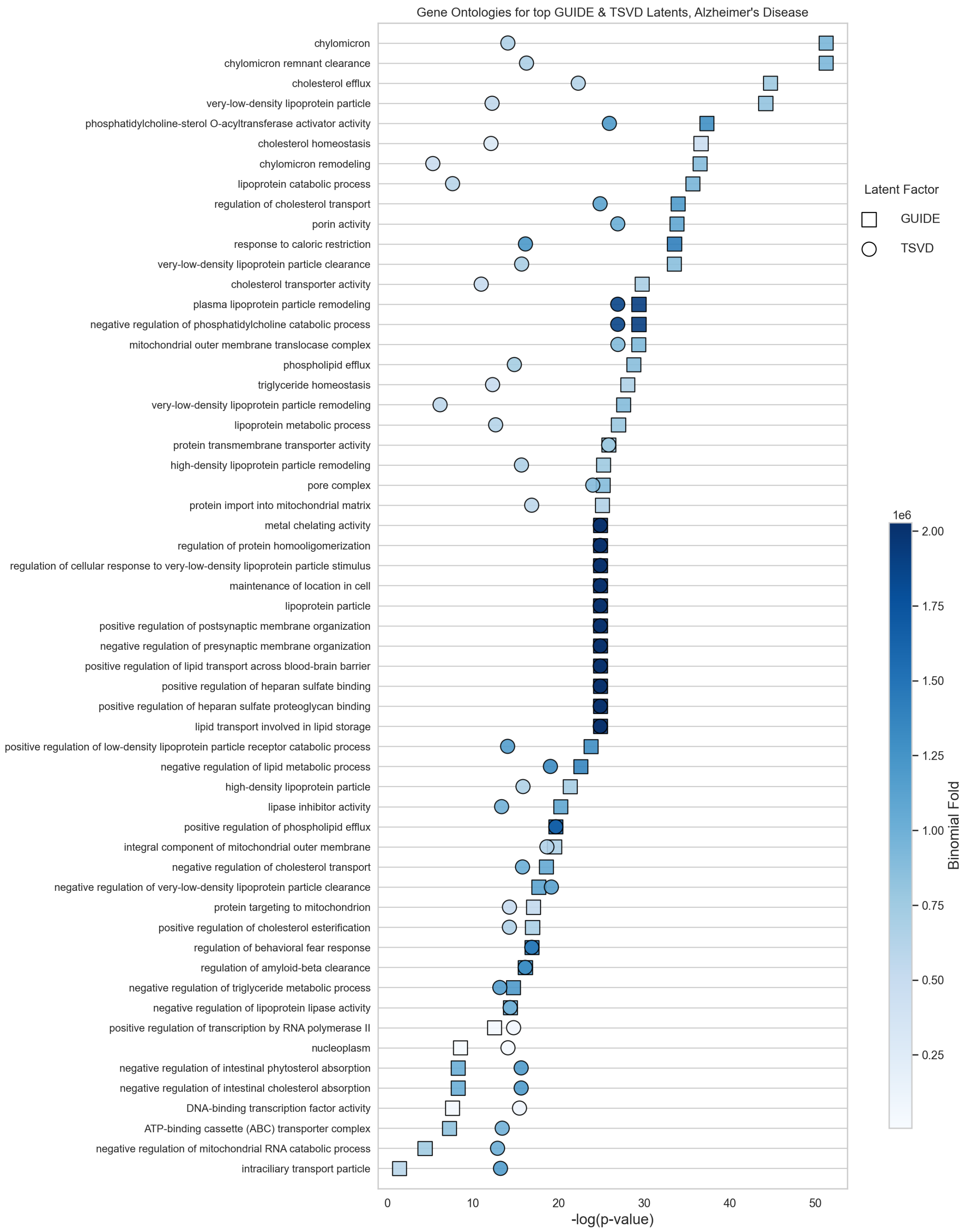


Figure 15: Gene Ontology (GO) Biological Processes for the top latent factors for Alzheimer's disease identified by each method, GUIDE latent 72 and TSVD latent 71. This factor is highly enriched for loci near the *APOE* genes, and exclusively prioritizes processes involved in cholesterol metabolism and processes implicated in Alzheimer's disease, in particular. Although this figure shows only shared processes, we note that TSVD latent 71 also prioritizes many processes that are not related to Alzheimer's pathophysiology (see main text).

Table 3: A group of 8 traits with the corresponding number of variants that are significant for each trait ($p < 5 \times 10^{-8}$) and the number of traits that are correlated to each trait with $r^2 > 0.5$

| Variants Significant For | No. Variants | No. Traits |
|---|---|---|
| BMI | 2123 | 335 |
| Ever Smoked | 137 | 212 |
| Standing Height | 12163 | 414 |
| Diabetes (diagnosed by doctor) | 1007 | 194 |
| Alcohol Intake Frequency | 520 | 121 |
| High Blood Pressure | 1183 | 344 |
| Miserableness | 403 | 66 |
| Seen Psychiatrist for Nerves, Anxiety, Tension or Depression | 376 | 60 |

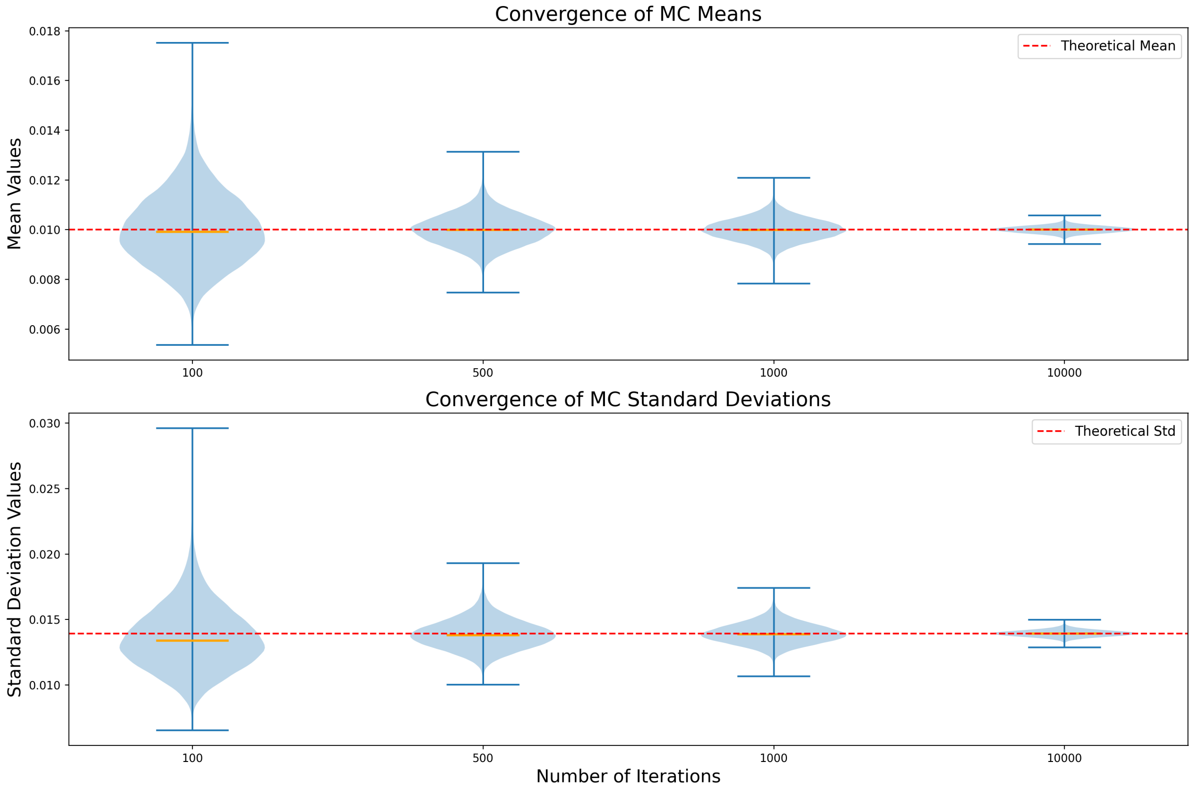


Figure 16: Violin plots comparing the means and standard deviations of $w$-values estimated using the Monte Carlo (MC) method of sampling independent points to estimate the null distribution versus the theoretical value given by the closed form solution, as described in Sec. 3.6 of the main text.

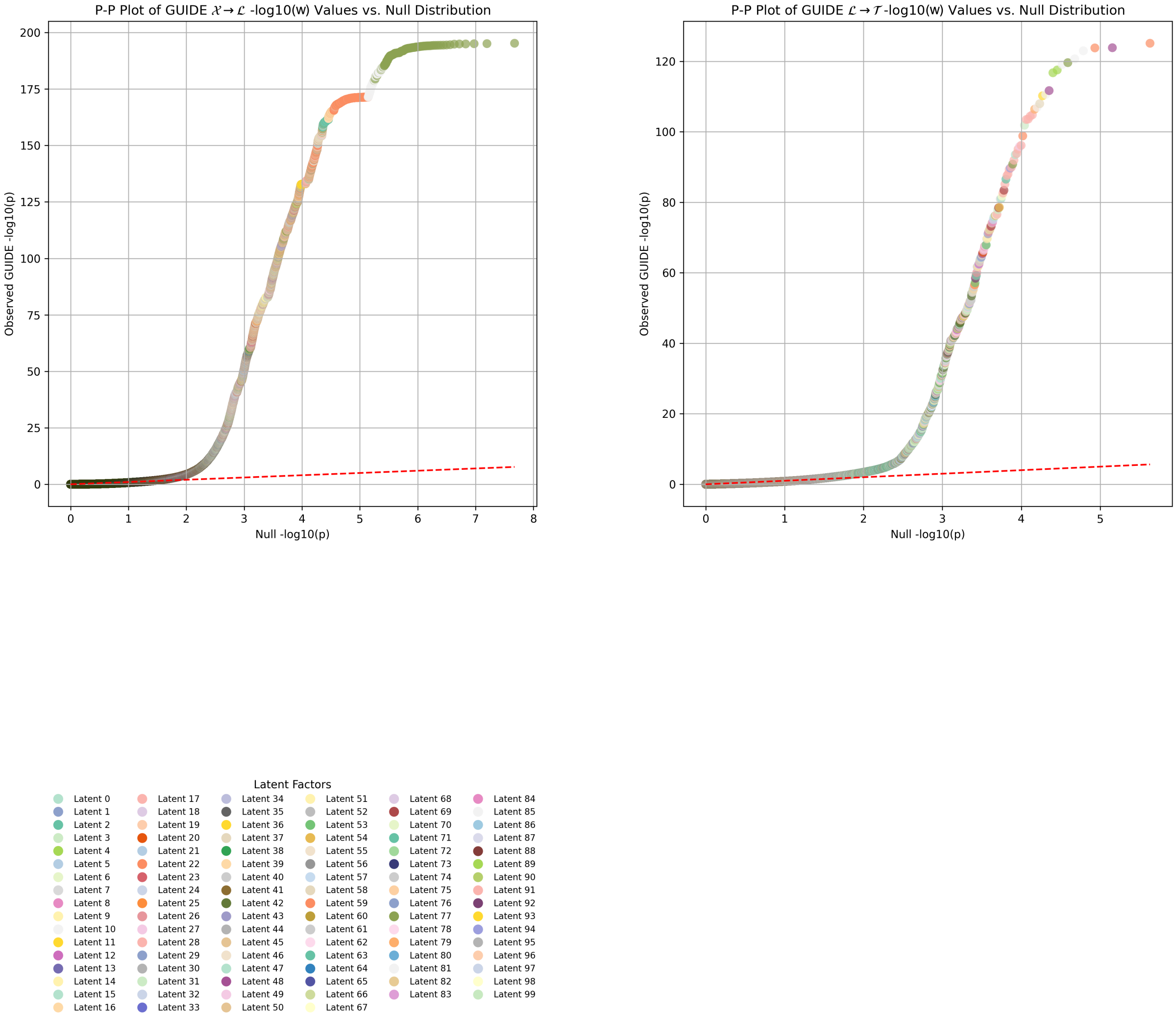


Figure 17: P-P plots of observed GUIDE $\mathcal{X} \rightarrow \mathcal{L}$ and $\mathcal{L} \rightarrow \mathcal{T}$ values versus the $-\log_{10}(w)$ values drawn from the theoretical null distribution, as described in Methods.

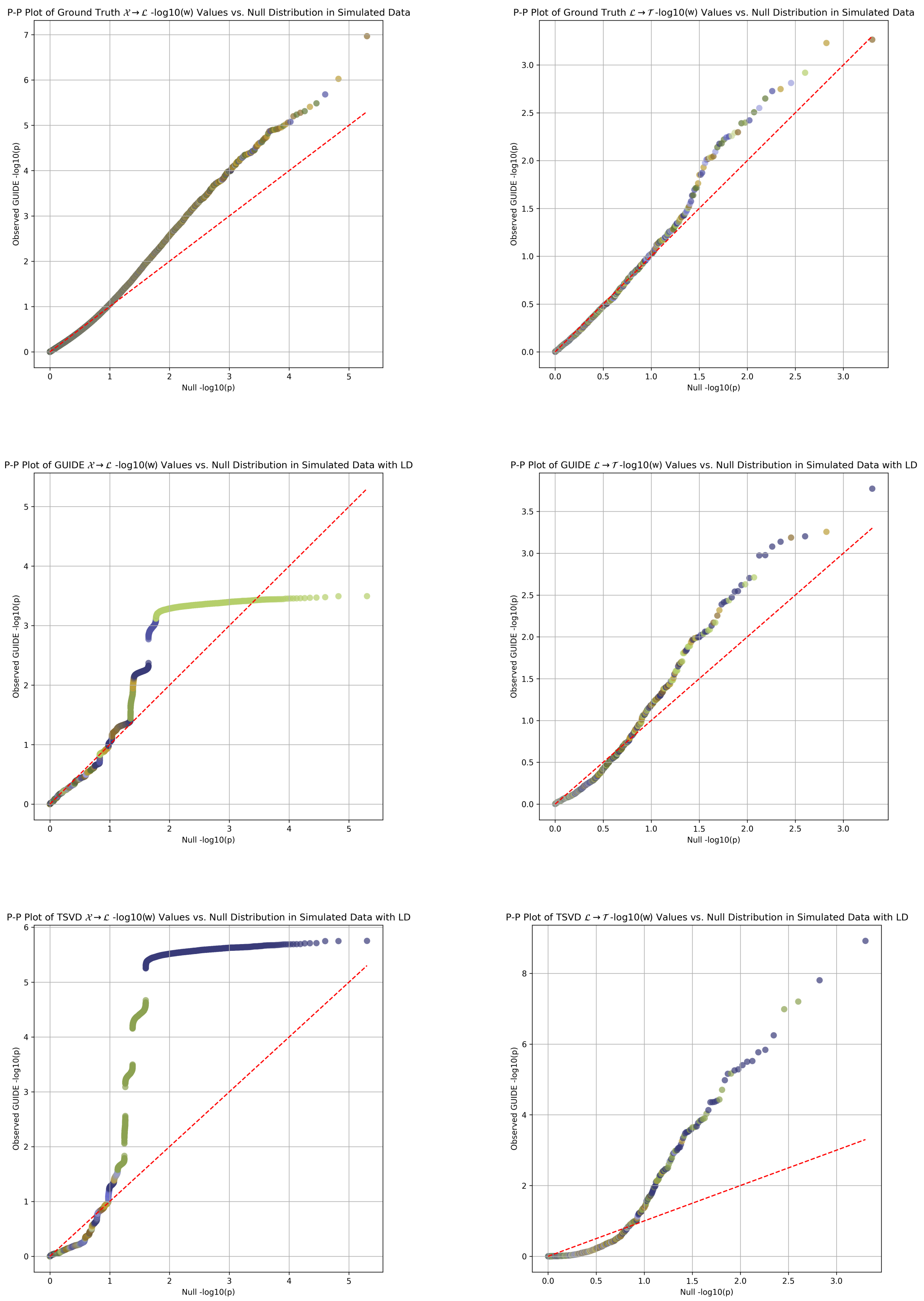


Figure 18: The "ground truth" plots are with the original simulated data without LD. The added LD for the rest of the plots is between 0 and 0.9. For these simulations, there were 10,000 variants, 100 traits and 10 latent factors. The legends are consistent for a given model's $\mathcal{X} \rightarrow \mathcal{L}$ and $\mathcal{L} \rightarrow \mathcal{T}$ plots but not between models.

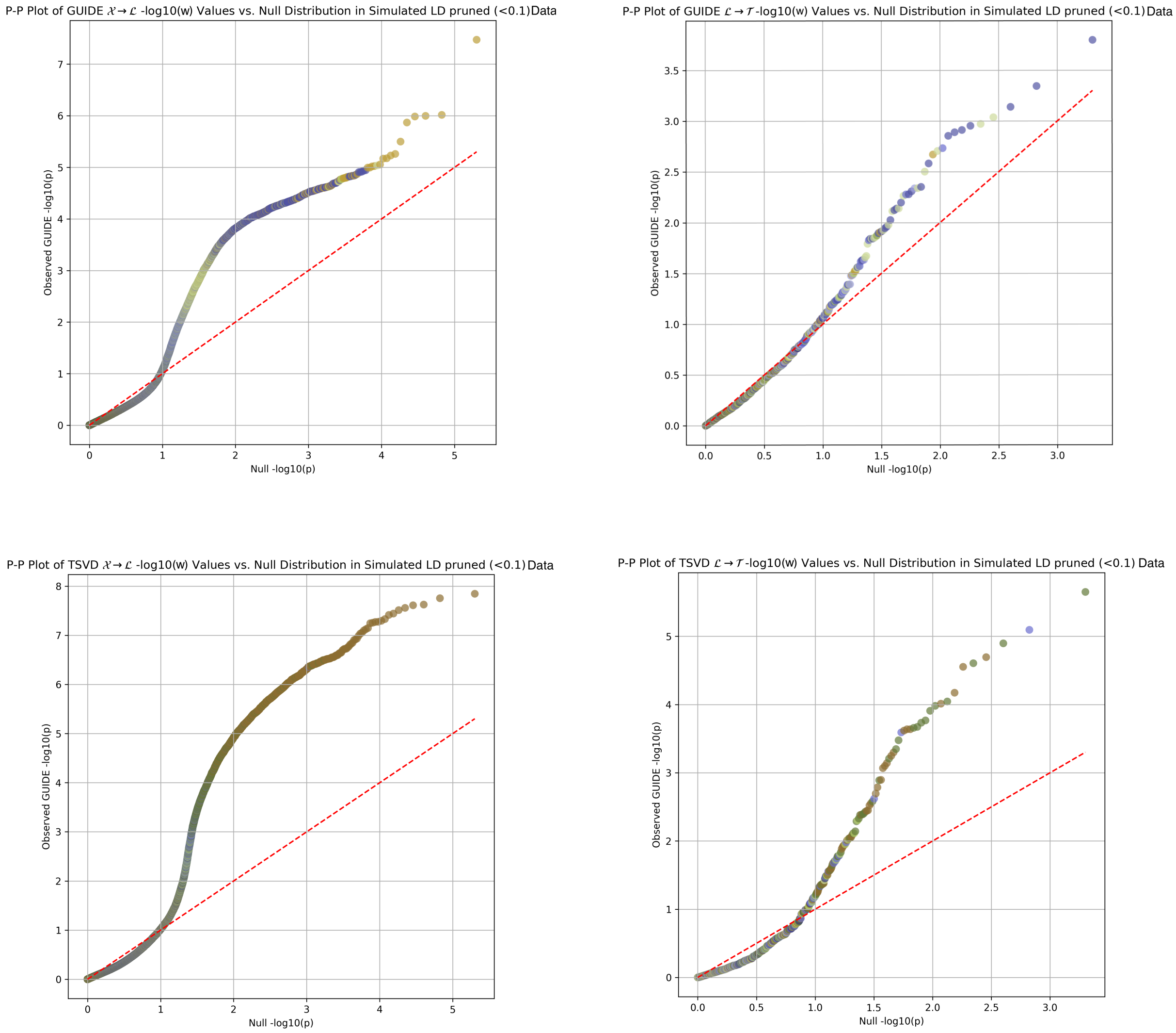


Figure 19: The same simulation as in Fig. 20 was used here, but with LD $< 0.1$.
